# DNA methylation-dependent and -independent binding of CDX2 directs activation of distinct developmental and homeostatic genes

**DOI:** 10.1101/2024.02.11.579850

**Authors:** Alireza Lorzadeh, George Ye, Sweta Sharma, Unmesh Jadhav

**Affiliations:** Department of Stem Cell Biology and Regenerative Medicine, Keck School of Medicine, University of Southern California, Los Angeles, CA 90033, USC; Norris Comprehensive Cancer Center, Keck School of Medicine, University of Southern California, Los Angeles, CA 90033, USC

## Abstract

Precise spatiotemporal and cell type-specific gene expression is essential for proper tissue development and function. Transcription factors (TFs) guide this process by binding to developmental stage-specific targets and establishing an appropriate enhancer landscape. In turn, DNA and chromatin modifications direct the genomic binding of TFs. However, how TFs navigate various chromatin features and selectively bind a small portion of the millions of possible genomic target loci is still not well understood. Here we show that Cdx2 - a pioneer TF that binds distinct targets in developing versus adult intestinal epithelial cells - has a preferential affinity for a non-canonical CpG-containing motif *in vivo*. A higher frequency of this motif at embryonic and fetal Cdx2 target loci and the specifically methylated state of the CpG during development allows selective Cdx2 binding and activation of developmental enhancers and linked genes. Conversely, demethylation at these enhancers prohibits ectopic Cdx2 binding in adult cells, where Cdx2 binds its canonical motif without a CpG. This differential Cdx2 binding allows for corecruitment of Ctcf and Hnf4, facilitating the establishment of intestinal superenhancers during development and enhancers mediating adult homeostatic functions, respectively. Induced gain of DNA methylation in the adult mouse epithelium or cultured cells causes ectopic recruitment of Cdx2 to the developmental target loci and facilitates cobinding of the partner TFs. Together, our results demonstrate that the differential CpG motif requirements for Cdx2 binding to developmental versus adult target sites allow it to navigate different DNA methylation profiles and activate cell type-specific genes at appropriate times.

**Graphical Abstract:** 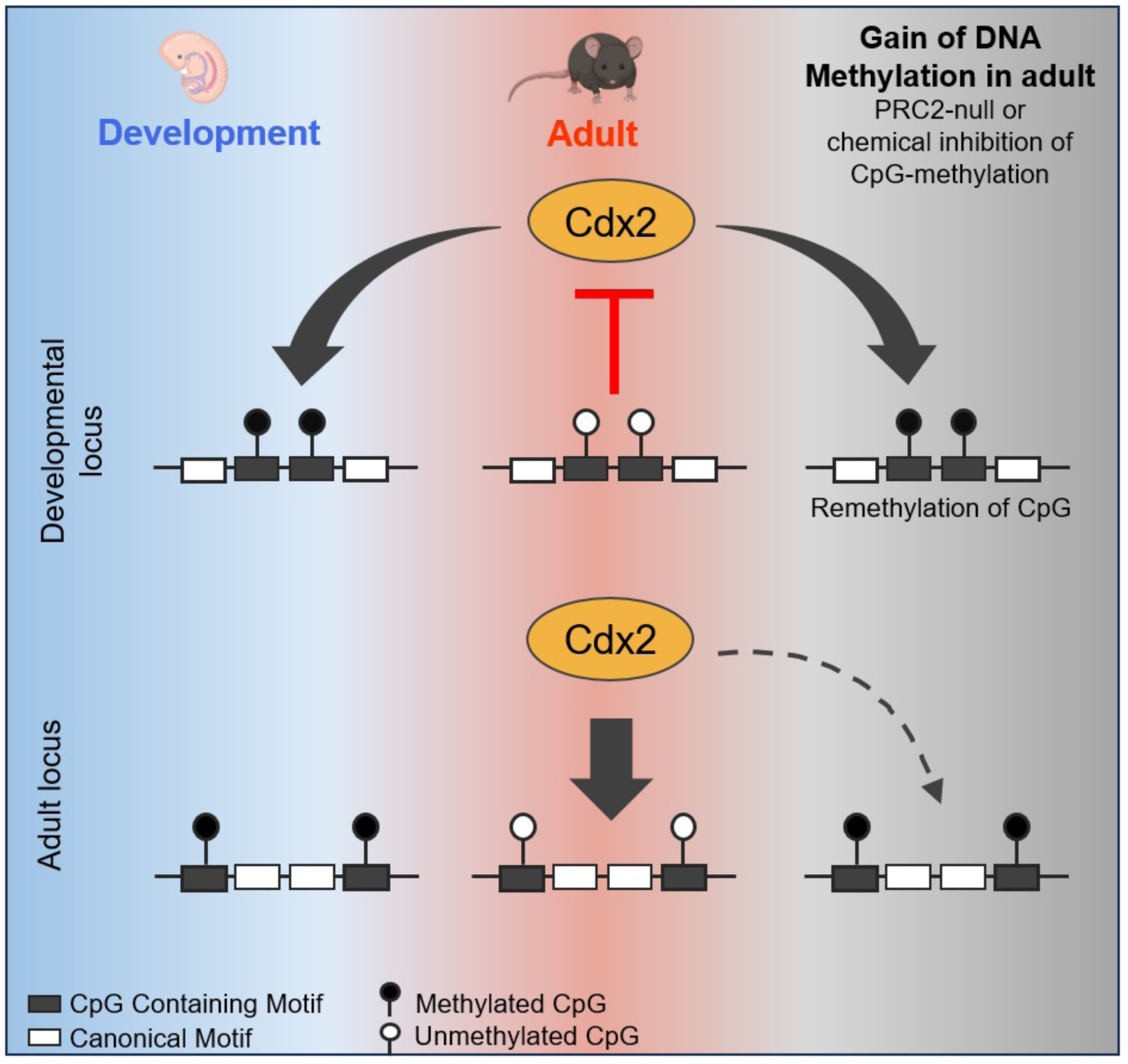

## INTRODUCTION

The establishment of functional cell types in adult tissues requires precise gene expression control throughout development. TFs recognize and bind to specific and short genomic sequences called target motifs within gene promoters and cis-regulatory enhancers giving rise to distinct transcriptional programs. Typically, TF motifs occur millions of times in the mammalian genome, however, most TFs only bind a fraction of those (D’haeseleer, 2006; Hansen et al., 2012). How TFs achieve such cell type-specific binding is not fully understood. Multiple factors including DNA and histone modifications, nucleosome positioning, chromatin compaction, and motif frequency and variation may play a role (Adams and Workman, 1995; Domcke et al., 2015; Isbel et al., 2022; Wunderlich and Mirny, 2009; Zaret, 2020). Pioneer TFs can overcome chromatin barriers to facilitate the recruitment of other TFs, formation and activation of gene regulatory enhancers, and gene transcription during tissue development and adult homeostasis (Slattery et al., 2014). To investigate such molecular controls *in vivo*, we utilized the intestinal epithelium as a model system. This single-cell thick inner lining of the intestine develops from a well-defined region of embryonic gut endoderm (Figure S1A). In adults, stem cells residing in the intestinal crypts continuously divide to produce terminally differentiated villus cells, which are predominantly absorptive enterocytes (Gehart and Clevers, 2019). Cdx2, a homeodomain-containing pioneer TF, is expressed in the intestinal epithelium starting from early embryonic development through adult life and controls specific gene targets during various stages of epithelial development and in adult cells. Accordingly, loss of Cdx2 action in early visceral endoderm leads to epithelial loss or squamous epithelial conversion (Chawengsaksophak et al., 1997; Gao et al., 2009; Tamai et al., 1999), while inactivation of Cdx2 in embryonic intestinal epithelium induces expression of gastric tissue markers (Grainger et al., 2010). On the other hand, Cdx2 loss in the mouse intestinal epithelium during late fetal development or in adult tissue impacts cellular functions linked to digestion and causes ectopic expression of genes from other parts of the digestive tract (Hryniuk et al., 2012; Simmini et al., 2014; Verzi et al., 2010; Verzi et al., 2011). Cdx2 achieves such developmental stage-specific gene control by binding to diverse genomic targets and establishing dynamic chromatin accessibility patterns across developing epithelium as well as in adult cells (Kumar et al., 2019). This system offers a unique opportunity to understand mechanisms that dynamically control TF interaction with epigenetic and chromatin barriers across the lifespan to set up cell type and developmental stage-specific enhancers and gene expression patterns.

Methylation at cytosine in the CpG dinucleotide is a well-studied epigenetic modification linked to transcriptional repression. Almost all CpGs across the vertebrate genome are methylated except for the ones at the active regulatory regions like promoters and enhancers (Bird et al., 1985; Schmitz et al., 2019). DNA methylation may further influence the formation and activity of these regulatory regions by impacting TF binding, which is central to establishing the open chromatin and facilitating epigenetic modifications. Recent high-throughput *in vitro* studies characterized the effect of CpG methylation on binding of more than 500 human TFs to various DNA binding domains using SELEX (Systematic Evolution of Ligands by Exponential Enrichment), showing that more TFs prefer methylated motifs (34%) as compared to the ones being inhibited (23%) when containing a CpG in their motif (Yin et al., 2017). These findings suggest that sensitivity to DNA methylation may be a major factor controlling TF binding *in vivo.* In addition to the CDX2 binding at the canonical motif without a CpG (ATAAA), CDX2 was shown to bind nucleotides with motifs containing CpGs with high affinity when the cytosine was methylated *in vitro* (Yin et al., 2017). However, for most TFs (including CDX2), it remains unknown if DNA methylation influences their genomic binding *in vivo*, where additional controlling factors may include DNA sequence surrounding the motif, nucleosome presence, and chromatin modifications. Our studies have previously revealed the genome-wide dynamics of DNA methylation in developing epithelial endoderm, where specific promoter and enhancer loci are demethylated stepwise to produce the adult chromatin landscape (Jadhav et al., 2019). Here we asked how Cdx2 navigates and impacts this chromatin landscape based on its capacity to bind variable motifs with and without DNA methylation and other chromatin controls. We demonstrate that Cdx2 binding across development and in adult cells is influenced by the distribution of its CpG-containing and canonical (not-CpG-containing) motifs. Prevalence of the CpG-containing motif at Cdx2’s developmental targets and methylation of the CpG within this motif facilitates Cdx2 binding during development and the absence of the CpG methylation at this motif serves to avoid ectopic Cdx2 recruitment at these loci in adult cells. Using two independent methods, we modulated DNA methylation through genetic elimination of Polycomb action in mouse interstitial epithelial cells and more directly using a chemical inhibitor in cultured cells. We show that induced methylation at the CpG-containing motifs causes ectopic Cdx2 recruitment at developmental targets in adult cells. Based on this CpG methylation dependent biding, Cdx2 facilitates recruitment of TFs Ctcf and Hnf4a to distinct genomic targets to establish developmental versus adult enhancer patterns and gene activation in the mammalian intestinal epithelium.

## RESULTS

### Distinct promoter- and enhancer-based gene control by Cdx2 in developing and adult intestinal epithelium

Pioneer factors such as Cdx2 overcome chromatin barriers including DNA and chromatin modifications to access their target motif sequences across the genome (Iwafuchi-Doi et al., 2016; Meers et al., 2019; Mingay et al., 2018). During mouse small intestinal development, where cell numbers are limiting, CUT&RUN assays allow precise detection of TF binding using just a few thousand cells as compared to the traditional ChIP-seq assays which require many times more cells as input (Meers et al., 2019). We used 50,000 purified intestinal epithelial cells to determine Cdx2 binding at embryonic day 12.5 (E12.5), fetal timepoint 16.5 (E16.5), and in adult differentiated villus cells (Figure S1A-B). CUT&RUN allowed us to identify a substantial number of Cdx2 binding sites in the developing epithelium (20,880 and 24,119 at E12.5 and E16.5, respectively) as well as adult cells (27,737). Importantly, a large number of Cdx2 binding events were specific to either development (7,806 in E12.5 and E16.5) or adult villus cells (8,901) (Figure 1A). Further classification of these loci based on 2-fold change in Cdx2 binding from E12.5 to E16.5 or E16.5 to adult (*q <* 0.01) divided the developmental and adult specific Cdx2 binding in 3 subgroups each (Dev 1-3 and Adult 1-3, respectively, Figure 1A, S1C). In line with previous findings (Kumar et al., 2019), 60% of all of the 8,901 adult binding sites (Adult 1-3) showed Cdx2 binding (MACS2 identified peak) at E16.5, a fetal timepoint representing advanced development, while only 8% of the Cdx2 binding at E12.5 (Dev 1-2) could be detected in adult cells (Figure 1A). This highlights the disparity in Cdx2 binding in developing versus adult intestinal epithelial cells and suggests a need for distinct underlying molecular controls for temporally selective Cdx2 recruitment. Genes linked to the developmental versus adult Cdx2 binding loci (Cdx2 peak +/- 25kb of the gene transcription start site, TSS) showed corresponding loss or gain of expression (Jadhav et al., 2019) (Figure S1D), suggesting a direct role of Cdx2 in regulating transcription of multiple stage-specific genes.

**Figure 1:**
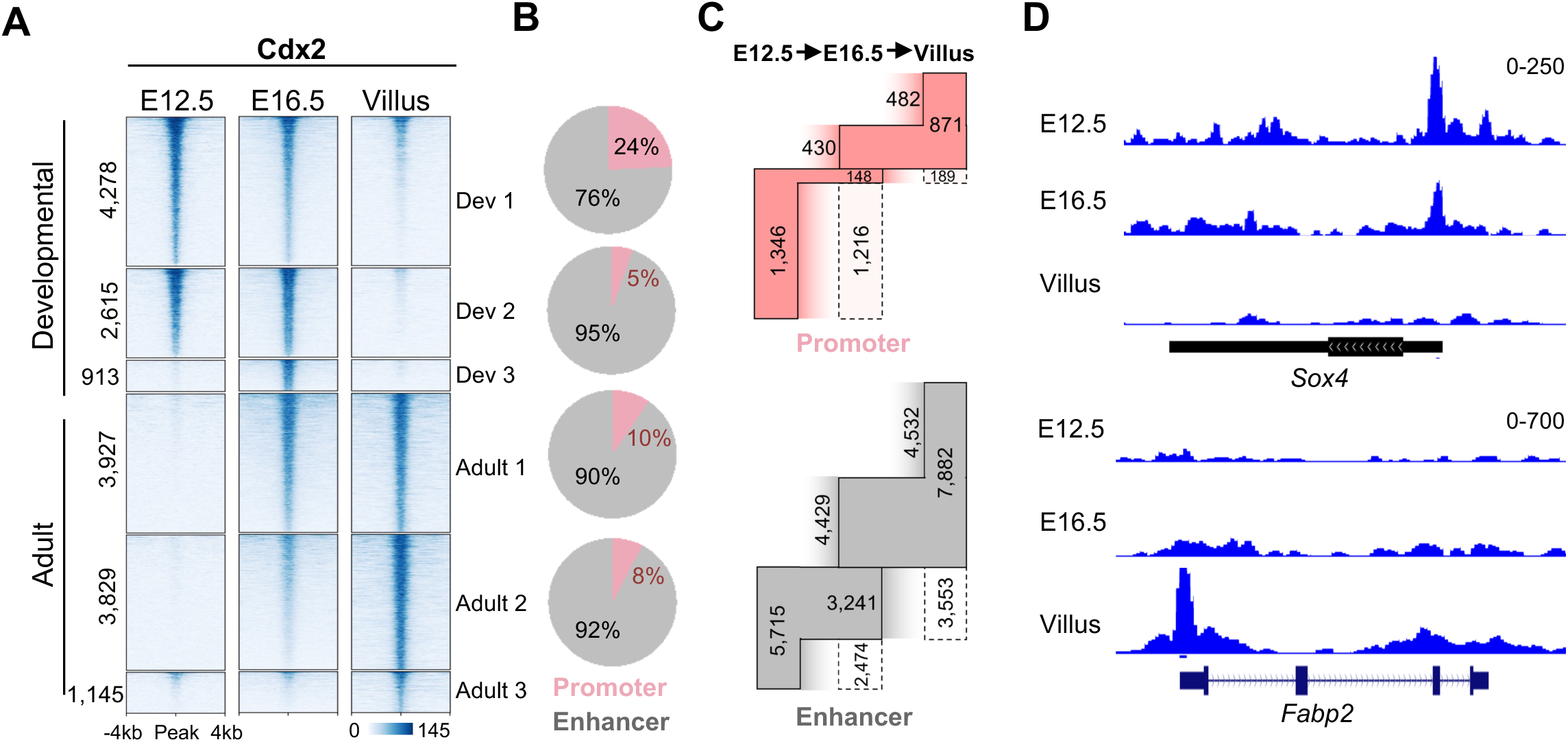
Evolution of Cdx2 binding across gene promoters and enhancers supports developmental and homeostatic functions in intestinal epithelium. **(A)** Heatmap showing Cdx2 CUT&RUN signal at its developmental and adult specific binding sites in embryonic (E12.5), fetal (E16.5) and adult epithelial (villus) cells; Dev 1-3 and Adult 1-3 subgroups represent 2-fold change (*q* < 0.01) in Cdx2 signal among the three timepoints. **(B)** Pie charts showing percentages of Cdx2 binding sites shown in (A) (Dev 1, 2 and Adult 1, 2) within and outside promoters (TSS -2 kb to +1 kb). **(C)** Gain, retention, and loss of Cdx2 binding at promoters (top panel) and enhancers (bottom panel) during development from E12.5 into adult epithelial cells; numbers outside the boxes and inside the dotted boxes represent gains and losses of Cdx2 binding at each of the timepoints, respectively. **(D)** Genome browser tracks showing progressive loss or recruitment of Cdx2 through epithelial development at *Sox4* and *Fabp2* gene promoters, respectively. See also Figure S1

To test if Cdx2 engages in distinct promoter- or enhancer-driven control of target genes in developing and adult epithelium, we classified the genomic loci with dynamic Cdx2 binding (Dev 1-3 and Adult 1-3, Figure 1A) based on their proximity to gene TSSs. At the loci with highly enriched Cdx2 occupancy during embryonic development (E12.5 Dev1, Figure 1A), 24% of the target genes had promoter based Cdx2 binding (2 kb upstream and 1 kb downstream of the TSS) as compared to all other groups (Dev 2-3, and Adult 1-3, Figure 1B and Figure S1E), which had average 8% and maximum of 13% promoter occupancy. On the other hand, majority (> 87%) of the Cdx2 binding sites in fetal (E16.5) and adult cells lie within regulatory regions outside the promoters. We confirmed that the temporal shift in Cdx2 binding from E12.5 going into adult cells resulted in altered chromatin accessibility and that the Cdx2 binding loci were bona fide promoter or enhancers by overlapping them with publicly available ATAC-seq data and active chromatin associated histone modification Histone 3 lysine 27 acetylation (H3K27ac, Figure S1F) (Jadhav et al., 2019; Kazakevych et al., 2017). The prevalent promoter-based Cdx2 binding in embryonic cells is quickly lost or reduced in fetal cells at E16.5 (90% of the 1,346 promoters at E12.5, Figure 1C-D), when significant morphological rearrangements, such as the formation of the crypt and villus structures, take place in the intestinal epithelium (Shyer et al., 2015; Walton et al., 2016). Conversely, Cdx2 binding is retained at 44% of the embryonic (E12.5) enhancers in fetal (E16.5) cells and 49% of this fetal binding is further retained in adult cells. Additionally, 65% and 58% of the enhancers in E16.5 and adult cells are newly bound by Cdx2 (Figure 1C, Figure S1G). This suggests a promoter-centric bias in gene targeting by Cdx2 in embryonic cells while transitioning into a more enhancer-driven control in adult cells during development. Importantly, the embryonic (E12.5) promoter targets of Cdx2 consist of genes critical for intestinal development such as *Sox4* and *Meis1* (Francis et al., 2019) (Figure 1D, Figure S1I). Interestingly, about 10% of promoter-based Cdx2 targets in Dev1-2 are other TFs, while only about 3% of the adult-specific Cdx2 binding is at TF promoters. This suggests that Cdx2 engages in directly activating other TFs critical for intestinal epithelial lineage commitment early on during development and engages in non-TF and functional gene activation through enhancers in adult tissue (Figure S1H).

### Pronounced presence of the CpG-containing Cdx2 motif at developmental binding sites

How various TFs depend on or overcome chromatin barriers to establish temporally distinct binding patterns remains poorly understood. In addition to the canonical binding motif containing the sequence ATAAA, studies show that Cdx2 binds to a CpG-containing motif with high affinity *in vitro* when the CpG (at the 4^th^ base in this motif) is methylated (Figure S2A). Based on these observations, we hypothesized that the differential Cdx2 sensitivity to distinct Cdx2 motifs may allow it to navigate and preferentially bind developmental or adult loci. To this end, we analyzed the Cdx2 binding dynamics across developmental and adult epithelial villus cells (Figure 1A) in the context of Cdx2 motif distribution and the temporal changes in CpG methylation. First, we determined the relative enrichment of the canonical and the CpG-containing motifs of Cdx2 (Figure S2A) at the Cdx2 target loci Dev 1-3 and Adult 1-3 (Figure 1A) using the motif enrichment analysis tool HOMER (Heinz et al., 2010). While the CpG-containing motif significantly enriched at the developmental loci (*p <* 10^-92^), it did not show enrichment at adult loci (Figure 2A). On the other hand, the canonical Cdx2 motif was present in both developmental and adult sites. As expected, the motif for Hnf4A was only enriched at adult loci, while motifs of Hoxc9 and Hoxd10 (with high sequence similarity to the CpG-containing Cdx2 motif), were enriched in developmental loci. To dissect the Cdx2 motif distribution across developmental and adult binding sites in detail, we used the FIMO algorithm which identifies significant instances of specific motifs within a set of input regions (Bailey et al., 2015). As seen in the HOMER analysis (Figure 2A), FIMO detected Cdx2 motifs (canonical or CpG-containing) more frequently at the developmental Cdx2 peaks in comparison to the adult loci, particularly near the center of the Cdx2 peaks (+/- 100 bp, *p*<0.005, Figure 2B); within 50 bp (+/-25 bp) from the center of the Cdx2 peaks, almost twice as many of the developmental loci had a detectable Cdx2 motif as compared to the adult loci (1,823 vs 986, Figure 2B).

**Figure 2.**
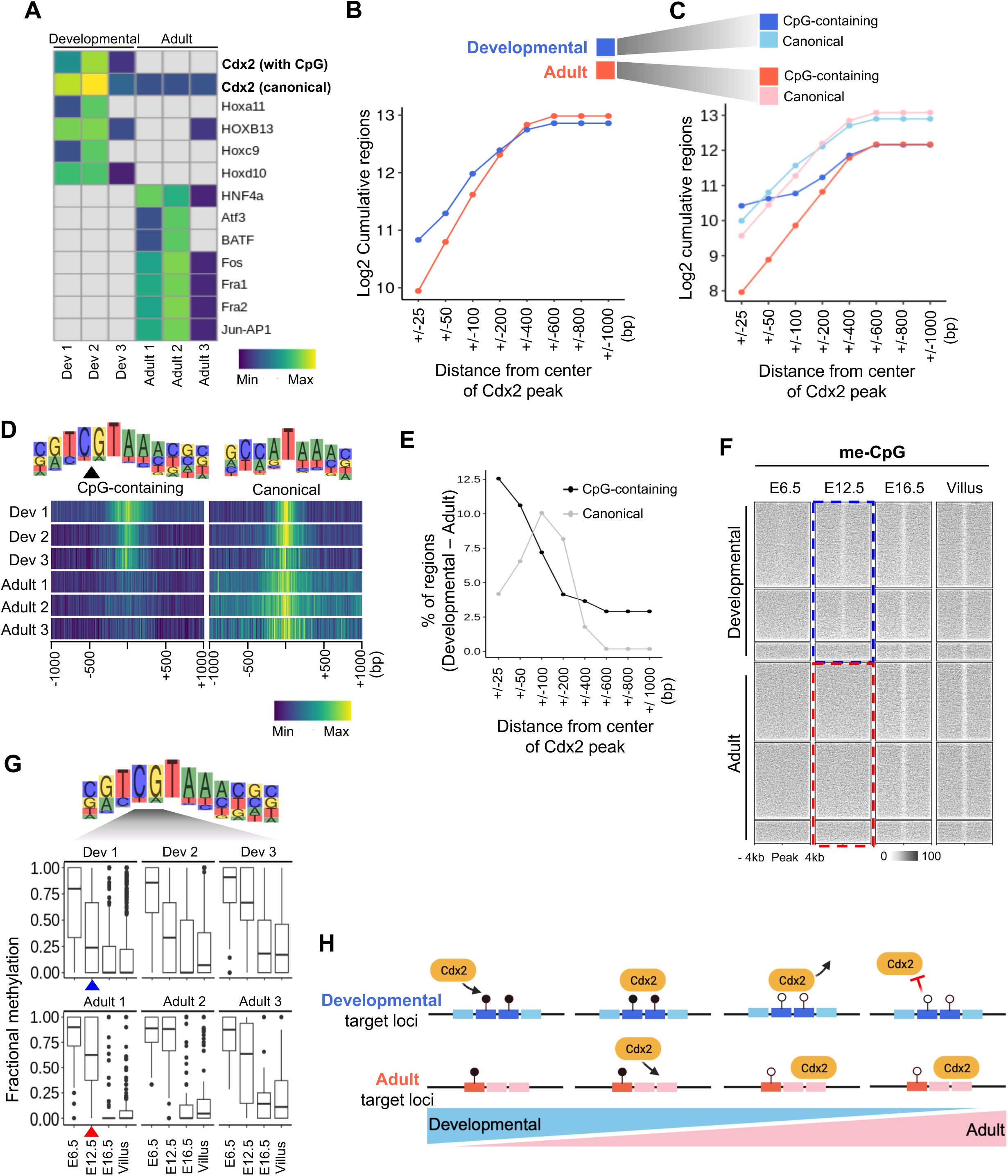
Heightened presence of CpG containing Cdx2 motif at its developmental binding sites. **(A)** Heatmap showing relative prevalence of TF motifs at developmental (Dev 1-3) and adult (Adult 1-3) cell specific Cdx2 binding sites as determined by Homer. **(B)** Line plot showing number of Cdx2 binding loci containing Cdx2 motifs within an increasing distance (indicated on the X-axis) from the center of the Cdx2 peak. **(C)** Line plot showing abundance of canonical and CpG-containing Cdx2 motifs at developmental (shades of blue) and adult (shades of red) cell specific Cdx2 binding sites within an increasing distance (indicated on the X-axis) from the center of the Cdx2 peak. **(D)** Density plots showing relative prevalence of CpG-containing motif near the center of developmental Cdx2 peaks (Dev 1-3) in comparison with the adult cell specific Cdx2 peaks (Adult 1-3); canonical Cdx2 motif shows widespread presence near Adult 1-3 sites in contrast with the Dev 1-3 loci. **(E)** Higher percent of regions with development specific Cdx2 binding contain CpG-containing motif as compared to regions with adult cell specific Cdx2 binding near the center of Cdx2 peak (within +/- 50 bp); beyond 50 bp of the peak center, canonical motif is more prevalent. **(F)** Heatmap showing DNA methylation at Cdx2 binding loci (as in Figure 1A) in early endoderm (E6.5) (Seisenberger et al., 2012), and embryonic (E12.5), fetal (E16.5), and adult epithelial (villus) cells (Jadhav et al., 2019); Cdx2 binding in embryonic epithelium is associated with early loss of DNA methylation (blue dotted box), while sites first bound by Cdx2 in the fetal epithelium show higher methylation levels (red dotted box). **(G)** Fractional methylation change at CpG (4^th^ base position) within the Cdx2 motif shows relatively early reduction in methylation at loci with developmental binding of Cdx2 (blue arrowheads) compared to binding in adult epithelium (red arrowheads). The CpG remains hypomethylated in adult villus at both developmental and adult loci. **(H)** A model for DNA methylation based Cdx2 binding at developmental target loci; requitement at adult target loci is driven by DNA methylation independent binding. While relative abundance of the CpG-containing motif and methylation at the CpG within the Cdx2 motif during development allows its early recruitment to these sites, demethylation at these loci protects them from ectopic Cdx2 binding in adult cells. See also Figure S2

We next compared the distribution of the canonical versus CpG-containing motifs in development and adult cell specific Cdx2 binding sites. While the canonical Cdx2 motif was present more frequently at both developmental and adult loci in comparison to the CpG-containing motif, developmental loci (Dev 1-3) showed higher prevalence of the CpG-containing motif in comparison to the adult loci (Adult 1-3), particularly within 50bp of the center of Cdx2 peaks (+/- 25bp, Figure 2C-D). In Dev 1 loci, where Cdx2 binds early during development, the CpG-containing motif was represented 14% more frequently than in Adult 1 loci, where Cdx2 is first recruited to establish a stable binding that lasts in the adult villus epithelium (Figure S2B). Overall, developmental sites with the CpG-containing motif exceeded the number of adult sites with this motif (12.5%, Figure 2E), suggesting that Cdx2 recruitment to distinct loci in embryonic and the adult epithelium may be linked to the relative prevalence of the different motifs. Higher prevalence of the CpG-containing motif towards the center of the developmental Cdx2 peaks and the relatively broader distribution of the canonical Cdx2 motif (Figure 2D, S2B) suggested a more focused targeting of Cdx2 at developmental loci. Indeed, the developmental Cdx2 peaks were significantly sharper (smaller genomic footprint) than the peaks in adult cells (Figure S2C). Interestingly, the developmental loci show more conservation across species as compared to the adult loci, suggesting a conserved molecular mechanism based on motif distribution and DNA methylation for Cdx2 recruitment to developmental targets (Figure S2D).

These data suggest that the greater prevalence of the CpG-containing Cdx2 motif near the center of developmental loci (Figure 2A-E) may selectively attract Cdx2 to these sites when the CpG in that motif is methylated, which happens only during development (Figure 2F). On the other hand, the canonical Cdx2 motif (non-CpG-containing, Figure S2A), which is more prevalent in adult Cdx2 target loci (Figure 2A-E) may underlie the recruitment of Cdx2 in adult cells. To test these possibilities, we looked at the DNA methylation dynamics associated with altered Cdx2 binding between developmental and adult timepoints (Figure 1A). Pioneer TFs such as NRF1 and PU.1 that preferentially bind methylated DNA can recruit demethylases like Tet2 to demethylate their binding site and surrounding CpGs, which creates an opportunity for other DNA methylation sensitive TFs to bind at the locus (de la Rica et al., 2013; Domcke et al., 2015; Terrell et al., 2023). Therefore, we considered the temporal changes in DNA methylation surrounding the Cdx2 binding peaks as well as at the CpG within the CpG-containing motif (4^th^ base position, Figure S2A). In early embryonic endoderm (E6.5) (Seisenberger et al., 2012), we see that all Cdx2 binding loci (Dev 1-3 and Adult 1-3) are fully methylated, including the CpG within the Cdx2 motif (Figure 2F-G). During early embryonic development of the epithelium, only the loci with early development specific binding of Cdx2 (Dev 1 and Dev 2) show average 23% and 32% drop in DNA methylation at E12.5 and methylation at the CpG within the Cdx2 motif is also reduced starting at E12.5. This is followed by further decrease in the DNA methylation at these loci by E16.5 (Dev 1: 30%, Dev 2: 48%, Dev 3: 49%) and in adult cells (Dev 1: 31%, Dev 2: 48%, and Dev 3: 50%, Figure 2F). This progressive loss of DNA methylation at developmental loci and the CpG within the Cdx2 motif is accompanied by loss of Cdx2 binding at these sites starting at E16.6 and into adult cells (Figure 1A), which suggests Cdx2’s dependence on the methylated DNA for its sustained binding at the developmental sites. This may explain how Cdx2 achieves a development-specific binding at these loci as the lack of DNA methylation at these sites in adult cells may preclude its binding (Figure 2G). On the other hand, the sites bound by Cdx2 specifically in adult cells (Adult 1-3) only show demethylation starting at E16.5, concordant with the beginning of Cdx2 binding (Figure 1A, 2F), and progressive loss of DNA methylation at these loci is accompanied by increasing Cdx2 binding at the canonical motif (without CpG) in adult cells in contrast to the developmental sites.

Alongside the dependence of Cdx2 on methylation at the CpG within its CpG-containing motif for developmental stage-specific binding, we considered the possibility that other chromatin features, particularly repressive modifications, may inhibit its binding at these loci in adult cells. In this regard, we conducted native-ChIP-seq, an assay particularly suitable for detecting the genome-wide distribution of histone modifications using low cell number input (Lorzadeh et al., 2016; Lorzadeh, 2017). DNA methylation and Histone 3 lysine 27 trimethylation (H3K27me3), a modification deposited by the polycomb repressive complex 2 (PRC2), are known to have opposing genome-wide distribution (Lorzadeh et al., 2021; Meehan and Pennings, 2017; Yu et al., 2019). DNA methylation and H3K27me3 form alternative modes of gene silencing as unmethylated genomic loci, particularly gene promoters, show widespread H3K27me3 deposition in mammalian cells (Jadhav et al., 2016; Xie et al., 2013; Yu et al., 2019). In ES cells and other *in vitro* systems, loss of DNA methylation or H3K27me3 causes alterations in the distribution of the other (Hagarman et al., 2013; McLaughlin et al., 2019; Reddington et al., 2013). Given this antagonistic relationship between DNA methylation and H3K27me3, we first looked at whether loss of DNA methylation at developmental loci in adult cells is accompanied by gain of H3K27me3. A small fraction of developmental sites (<10%) show deposition of H3K27me3 in adult cells (Figure S2E), suggesting no significant association with or impact on Cdx2 binding. H3K9me3, a modification associated with chromatin compaction, and H2AK119Ub, which is linked to active as well as inactive chromatin loci, showed very little concordance with Cdx2 binding in adult cells (Figure S2E). Interestingly, adult cells show H3K36me2, a histone modification known to recruit the de novo DNA methylase DNMT3A, at the developmental loci despite the hypomethylated state of the enhancers (Weinberg et al., 2019) (Figure S2E).

Together, these data strongly suggest that the genomic targeting of Cdx2 in developing and adult intestinal epithelial cells is based on variability in its target motif sequence, motif distribution, and sensitivity to DNA methylation. While methylation of the CpG within the CpG-containing motif may help recruit Cdx2 to developmental loci, the demethylation of this CpG post development may deter ectopic Cdx2 binding at these sites in the adult cells (Figure 2H).

### Progressive activation of enhancers by Cdx2 through clustered binding and corecruitment of Ctcf and Hnf4

Given the gradual increase of enhancer-based Cdx2 binding through development (Figure 1C), we examined the grouping of its binding and asked if Cdx2 facilitates corecruitment of TFs with different sensitivities to chromatin features. For the development and adult cell specific target sites of Cdx2, (Dev 1-3 and Adult 1-3, Figure 1A), we asked if there were other Cdx2 binding instances within 50 kb. While 60% of the embryonic sites (Dev 1 and Dev 2) had no other Cdx2 binding within 50 kb, 71% of the adult bound sites (Adult 1 and Adult 2) had at least one other site nearby (Figure 3A), and this clustering was apparent when considering Cdx2 binding only at the enhancers or when considering Cdx2 peaks only within 10 kb distance from each other (Figure S3A-B). Cdx2 binding around genes expressed in adult homeostatic cells such as *Krt19* was accompanied by significantly more clustering of Cdx2 in adult villus cells when compared to developmental sites (groups of median 3 Cdx2 peaks vs 2 peaks at adult and developmental loci, respectively, Figure 3C). Accordingly, 6,492 adult loci had 2 or more Cdx2 binding events surrounding them as compared to 3,283 developmental loci (*p* < 0.001). In combination with the increased enhancer-based binding of Cdx2 in the adult cells (Figure 1C), these data suggest that Cdx2 binding in groups or clusters of target loci drives the establishment of adult enhancer patterns.

**Figure 3.**
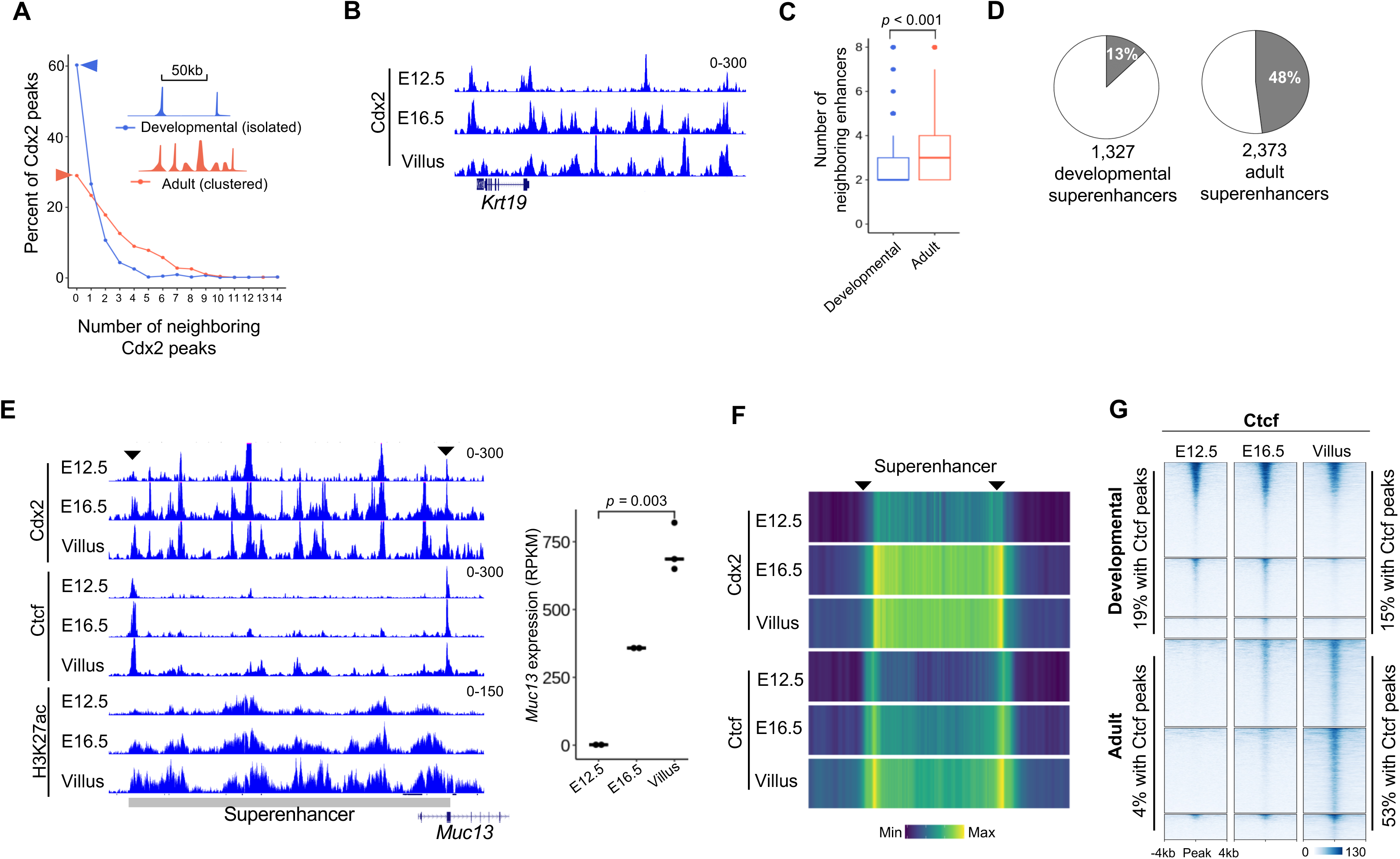
Cdx2 facilitates establishment of adult homeostatic superenhancers by directing Ctcf recruitment. **(A)** Percentage of developmental (blue) or adult (red) cell specific Cdx2 bound loci (as in Figure 1A) with designated number of other Cdx2 peaks nearby (+/- 50 kb); drawing in the inset represents isolated Cdx2 peaks (not within 50 kb of each other) in developing cells, while adult cells show multiple neighboring peaks in the same area; large number of (70%) of dynamic Cdx2 binding events that are solitary in developing epithelium (blue arrowhead) as opposed to only 40% in adult cells (red arrowhead). **(B)** Representative genomic tracks showing clustered binding of Cdx2 that grows through development near gene *Krt19*. **(C)** boxplot to the right shows number of neighboring (+/- 50 kb) dynamic Cdx2 bound enhancers (sites in Figure 1A) surrounding developmental (blue) or adult (red) cell specific Cdx2 binding loci. **(D)** Percentage of 1,327 developmental (E12.5 and E14.5) and 2,373 adult superenhancers that have dynamic Cdx2 binding during epithelial maturation. **(E)** Genome browser view of Cdx2, Ctcf, and H3K27ac signals at adult specific Cdx2 cluster at a superenhancers near *Mcu13* gene; black arrowheads indicate the superenhancer borders with Cdx2 and Ctcf cobinding. Plot on the right shows gain in expression of *Muc13* through development. **(F)** Heatmap showing Cdx2 and Ctcf signals at superenhancers in E12.5, E16.5 and adult villus. Superenhancers boundaries are shown with black arrowheads. **(G)** Heatmap showing Ctcf signal at developmental and adult specific Cdx2 binding sites (as in Figure 1A) in E12.5, E16.5 and adult villus cells; the regions are displayed in decreasing order of Ctcf signal. 19% and 15% loci with development specific Cdx2 binding show Ctcf occupancy at E12.5 and adult villus samples, respectively. 4% and 53% loci with adult specific Cdx2 binding show Ctcf binding in E12.5 and adult villus samples, respectively. See also Figure S3

Superenhancers are known to be hubs for coordinate binding of multiple TFs that facilitate wide-scale deposition of active enhancer mark H3K27ac to induce linked gene expression (Feng et al., 2023; Jia et al., 2020; Whyte et al., 2013). Using published data, we identified superenhancers at developmental timepoints (1,327 superenhancers combined at E12.5 and E14.5) and adult villus cells (2,373, superenhancers) (Jadhav et al., 2019; Kazakevych et al., 2017). While 13% of the developmental Cdx2 binding sites (Dev 1-3, Figure 1A) were located within superenhancers, 48% of the loci with adult cell specific Cdx2 binding (Adult 1-3, Figure 1A) overlapped with superenhancers (Figure 3D). Cdx2 showed binding at many of the superenhancer boundaries at E12.5, and the binding progressively spread into the body of superenhancers during development (Figure 3E-F), suggesting a role of Cdx2 in defining adult superenhancer borders through binding in early development.

Ctcf is a TF involved in chromatin organization and functions to form topologically associated domains (TADs) for genome compartmentalization and promoter-enhancer interactions using cohesion-based DNA looping. Various chromatin controls of Ctcf recruitment are being discovered, which include sensitivity to DNA methylation at CpGs within its motif (Figure S3C), and recruitment to its genomic targets through interaction with other TFs (Holwerda and de Laat, 2013; Liu et al., 2023; Maurano et al., 2015; Nanavaty et al., 2020; Ong and Corces, 2014; Yin et al., 2017). As Ctcf is often bound at the superenhancer boundaries (Islam et al., 2023), we conducted CUT&RUN assays to determine the dynamics of Ctcf recruitment during intestinal development and its dependance on Cdx2 based corecruitment. While only 4% of the adult cell specific Cdx2 sites (Adult 1-3) had Ctcf binding at E12.5, 53% of these loci gained Ctcf binding during development (Figure 3G). On the other hand, about 19% percent of the developmental Cdx2 target sites at E12.5 had Ctcf co-binding and 77% of this binding was maintained in adult cells with 15% of all developmental sites (Dev 1-3) showing co-binding with Cdx2 (Figure 3G). These data show a more prevalent interaction between Ctcf and Cdx2 at adult enhancers that temporally grows through development as compared to a limited cooperation at early developmental target sites starting at E12.5, which remains stable in the adult cells. Moreover, Ctcf binding at superenhancer boundaries follows Cdx2 binding temporally as it starts from either or both sides of the superenhancers in embryonic cells (E12.5) and spreads across the body of superenhancers (Figure 3E-F, S3D). This suggests that Cdx2 binding may be required to prime these loci for Ctcf binding in order to establish the adult superenhancers during development. On the other hand, when we conducted CUT&RUN for Hnf4, an intestinal TF important for homeostatic gene activation in the adult tissue, it showed high cobinding with Cdx2 only at the adult villus specific Cdx2 binding sites (Figure S3E). Moreover, at superenhancers, Hnf4 shows high signal through the body of the superenhancers (Figure S3F-G) in contrast to the stronger Ctcf binding at the boundaries (Figure 3E-F, S3D-F). Thus, temporal dynamics of Cdx2 binding in intestinal epithelium may facilitate step-wise establishment of developmental and adult enhancers through systematic corecruitment of two different TFs, Ctcf and Hnf4a, respectively.

### Loss of PRC2 action leads to altered DNA methylation and Cdx2 binding

Although the propensity of Cdx2 to bind the methylated CpG-containing motif has been noted by *in vitro* studies (Yin et al., 2017), its relevance to Cdx2 recruitment in cells and functional implications for gene expression or cellular identity remain unknown. To test this critically, we used two distinct experimental systems, a mouse model and a cell line, in which we determined if induced DNA methylation at the CpG-containing Cdx2 motifs in adult cells could promote ectopic recruitment of Cdx2 to its development-specific target sites. In mammalian tissues, many developmental genes lose DNA methylation at promoters when they are expressed during embryonic growth and H3K27me3 deposition in adult cells protects these genes from ectopic expression in homeostatic conditions (Jadhav et al., 2016; Xie et al., 2013). We and others have shown that genetic deletion of Ezh1/2, the enzymatic subunits of the PRC2 complex, or Eed, a protein that stabilizes the complex, cause loss of H3K27me3 in various adult tissues including intestinal epithelium, leading to reactivation of developmental genes (Ezhkova et al., 2009; Jadhav et al., 2016; Lee et al., 2015; Lien et al., 2011; Xie et al., 2013). Accordingly, we have previously used induced deletion of *Eed* in the entire mouse intestinal epithelium or specifically in the intestinal stem cells (ISCs) and characterized the temporal impact of H3K27me3 loss on promoter and enhancer activity and gene expression (Jadhav et al., 2019; Jadhav et al., 2020; Jadhav et al., 2016). Notably, loss of *Eed* in the intestinal epithelium caused activation of enhancers marked by hypomethylated DNA as they gained enhancer associated histone modifications H3K4me1 and H3K27ac, further leading to expression of linked genes. Using this well-established *in vivo* system for enhancer modulation, we asked how altered chromatin features, particularly DNA methylation, may influence Cdx2 binding.

We deleted *Eed* across the adult intestinal epithelium using 5 intraperitoneal injections of tamoxifen (1mg/dose) on consecutive days in *Eed^Fl/Fl^; Villin-Cre*^ER-T2^ mice as before (el Marjou et al., 2004; Jadhav et al., 2019; Jadhav et al., 2016) (Figure 4A) and and analyzed the genome-wide changes in DNA methylation using whole genome bisulfite sequencing (WGBS). To identify changes in DNA methylation in the *Eed^-/-^* cells, we conducted whole genome bisulfite sequencing (WGBS) and observed significant gains of DNA methylation at enhancers (Figure 4B). By comparing the methylation at 16,392,619 consensus CpGs between *WT* and the *Eed^-/-^* cells, we determined differentially methylated regions (DMRs, minimum 40% change in DNA methylation and *q* < 0.001); 82% of the DMRs (23,978 sites) showed gain of DNA methylation. Importantly, 95% of these DNA methylation gains were at enhancers, while 1,222 promoters showed hypermethylation in *Eed^-/-^* cells (Figure 4B). In a parallel approach, in *WT* cells we identified unmethylated regions (UMRs, with CpG methylation <10%), which mostly represent gene promoters, and low methylated regions (LMRs, with CpG methylation > 10% and < 60%) that represent enhancers (Burger et al., 2013; Jadhav et al., 2019; Stadler et al., 2011). While many promoters maintained an unmethylated state upon loss of *Eed* (UMRs), we observed a gain of methylation at many enhancers (LMRs) (Figure 4C). Additionally, we divided the genome into different states based on various chromatin features including activating and repressive histone modifications and TF binding using CHROMHMM algorithm (Ernst and Kellis, 2012). This revealed a gain of DNA methylation at enhancers in *Eed^-/-^* cells, particularly ones with Cdx2 binding (Figure 4D, Figure S4A-B). Together these data show that the gain of DNA methylation upon loss of *Eed* is focused at hypomethylated adult enhancers rather than a global hypermethylation of the genome. The gain of DNA methylation at enhancers was apparent at developmental Cdx2 binding loci (Dev 1-3), recapitulating a developmental CpG methylation state in adult cells (Figure 4E, Figure S4C). To examine the effect of this remethylation at developmental enhancer loci on Cdx2 binding, we conducted CUT&RUN for Cdx2 using adult *Eed^-/-^* villus cells. Cdx2 binding at developmental loci (Dev 1-3) was increased in *Eed^-/-^* villus cells (Figure 4F, G). Comparison of Cdx2 binding in *Eed^-/-^* cells with that in *WT* cells identified 1,053 Cdx2 binding events unique to *Eed^-/-^*cells (*q* < 0.01, 1.5X gain, Figure 5A). A significant number of these Cdx2 gains in *Eed^-/-^* overlapped with developmental sites (74%, hypergeometric *p*-value < 0.001) and the Cdx2 signal at these loci in *Eed^-/-^*cells was equivalent to that in the embryonic cells (E12.5) suggesting reinstatement of the developmental binding of Cdx2 (Figure 5B). Gene ontology analysis showed that the sites with Cdx2 recruitment in *Eed^-/-^* cells were associated with developmental genes regulating tissue patterning, further underscoring the reestablishment of developmental Cdx2 binding patterns in adult cells. Contrary to this, site with reduced Cdx2 binding were associated with genes involved in intestinal homeostatic functions (Figure S5A). Moreover, the DNA methylation level in *Eed^-/-^* cells at these sites was reminiscent of embryonic methylation level (E12.5) (Figure 5C, Figure S5B). We could detect the CpG-containing motif at twice as many loci (37%) with Cdx2 gain in the *Eed^-/-^* cells as compared to sites with reduced Cdx2 binding (17%), and the CpG at the 4^th^ base position in the Cdx2 motif showed gain of methylation (Figure 5D). Thus, our results strongly argue that the demethylation at developmental Cdx2 binding sites in adult cells may be sufficient to avoid its binding to these sites post development, and replenishing the embryonic DNA methylation state in adult cells can recruit Cdx2 to the developmental enhancers in adult epithelial cells. Importantly, the Cdx2 binding is necessary for open chromatin status at the developmental loci, as we see loss of ATAC-seq signal at these loci in adult cells, while Cdx2 binding in adult *Eed*^-/-^ cells results in re-opening the chromatin at these sites (Figure 5E-F), highlighting Cdx2’s pioneer factor function. The central role of Cdx2 in establishing developmental enhancers is further corroborated by the gain of H3K27ac signal indicating ectopic activation of these enhancers in adult *Eed*^-/-^ cells (Figure 5G) (Kazakevych et al., 2017).

**Figure 4.**
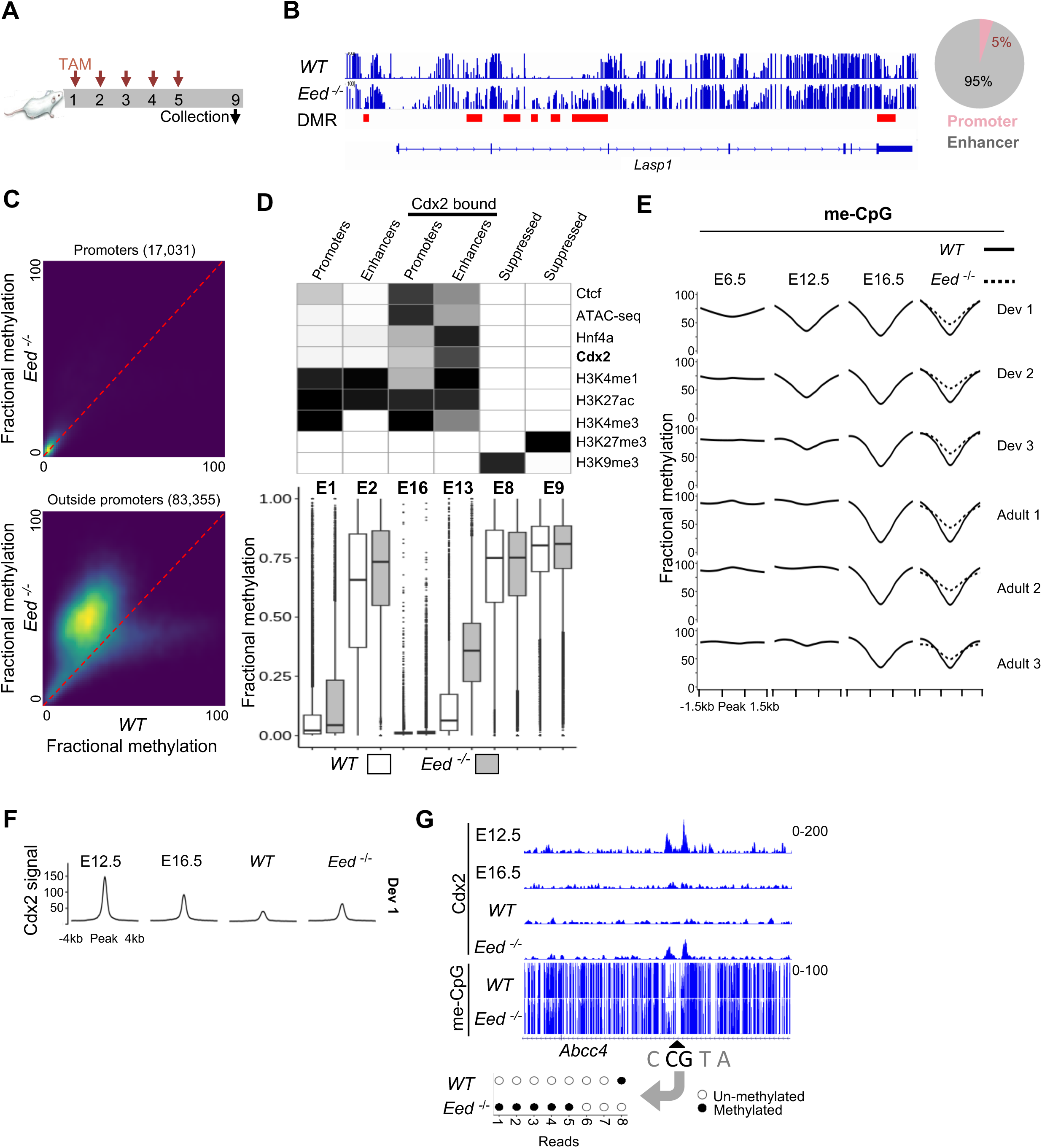
Loss of PRC2 activity causes gain of DNA methylation at enhancers leading to Cdx2 recruitment. **(A)** Experimental schematic showing 5 intraperitoneal injections of Tamoxifen (TAM) on consecutive days cause deletion of *Eed* across the intestinal epithelium; epithelial cells are collected 4 days after the last injection (experimental day 9). **(B)** Genome browser view showing gain of CpG methylation in *Eed^-/-^* epithelium at multiple enhancer loci near *Lasp1* gene that are identified as DMRs; promoter region remains unmethylated. Pie chart showing percentages of DNA methylation gains at promoters and enhancers. **(C)** Density plot showing DNA methylation at promoter (top panel) and non-promoter (bottom panel) linked UMRs and LMRs; LMRs (mostly representing enhancers) have the most prevalent gain of methylation in *Eed^-/-^* cells. **(D)** Heatmap showing genomic states identified by CHROMHMM based on TF binding and histone modifications corresponding to promoter, enhancer, and repressed chromatin; Cdx2 occupied enhancers (H3K4me1+ and H3K27ac+) particularly show gain in DNA methylation in *Eed^-/-^* cells, while promoters (H3K4me3+) and repressed chromatin regions (H3K27me3+ or H3K9me3+) show relatively stable unmethylated and methylated status, respectively. **(E)** DNA methylation profiles at development and adult specific Cdx2 binding sites (Dev 1-3 and Adult 1-3, respectively) showing the gain of methylation upon loss of PRC2 activity (*Eed^-/-^*). **(F)** Profile plot of Cdx2 signal show ectopic recruitment of Cdx2 at developmental loci (Dev 1) in adult *Eed^-/-^* cells in concordance with the gain of DNA methylation in (E). **(G)** Genomic tracks showing *Abcc4* gene locus with CpG-containing Cdx2 motif bound by the TF only during development, where gain of methylation at the Cdx2 binding locus in *Eed^-/-^* leads to recruitment of the TF; dot plot at the bottom shows methylated (solid black dots) or unmethylated (white dots) status of the CpG in Cdx2 motif (4^th^ position) in 8 independent sequencing reads from *WT* and *Eed^-/-^* WGBS data. See also Figure S4

**Figure 5.**
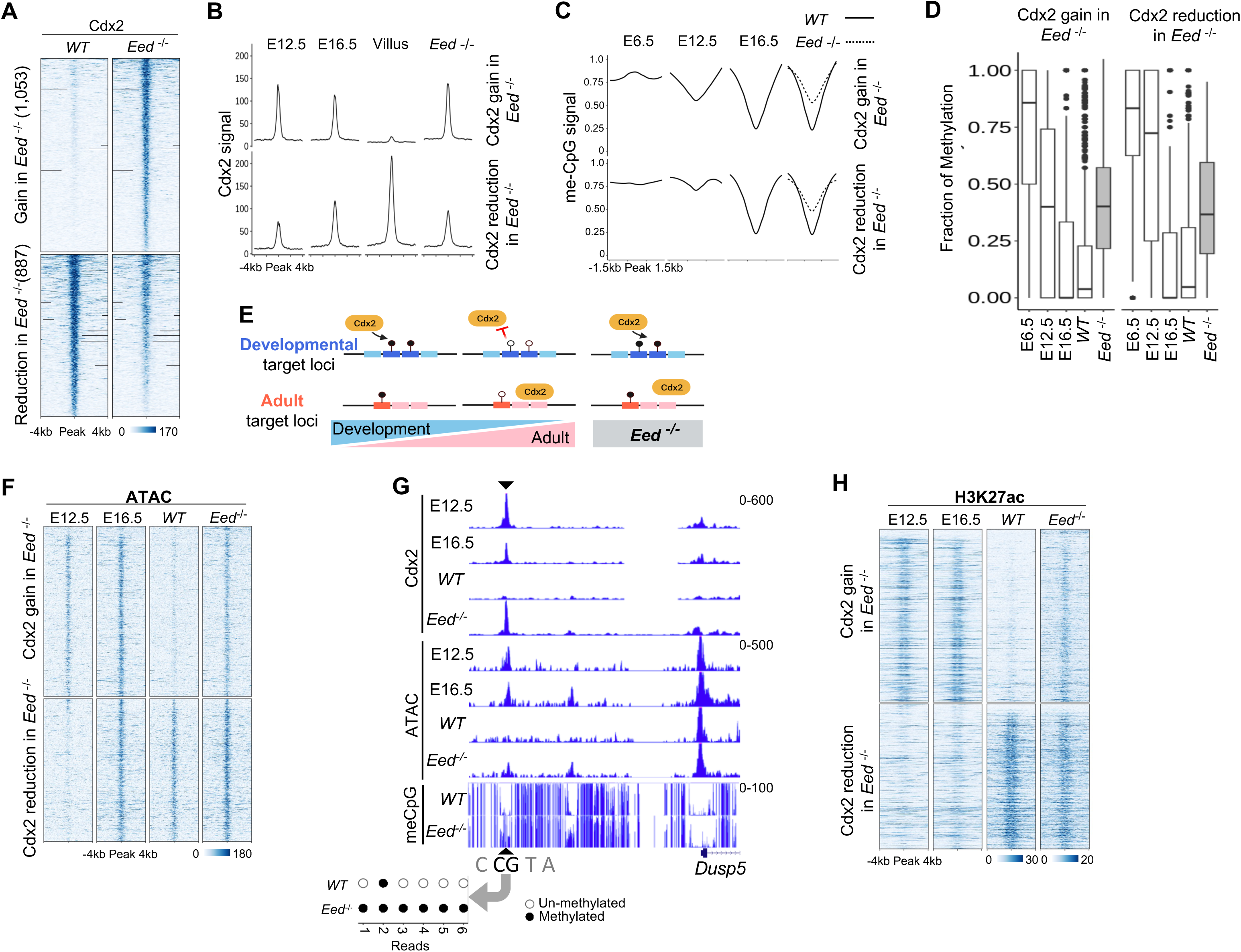
Altered Cdx2 binding in adult cells revives developmental Ctcf recruitment. **(A)** Heatmap showing significant (*q* < 0.05, 1.5X) gain or reduction of Cdx2 binding in *Eed*^-/-^ cells as compared to the native villus cells (*WT*). **(B)** Profile plots showing Cdx2 signal at loci with gain and loss of Cdx2 binding in *Eed*^-/-^ cells from embryonic (E12.5), fetal (E16.5), adult villus, and *Eed^-/-^*epithelium. **(C)** Profile plots showing DNA methylation signal at loci with gain or reduction of Cdx2 binding in *Eed*^-/-^ cells from embryonic (E6.5 and E12.5), fetal (E16.5), adult villus, and *Eed*^-/-^ epithelium. **(D)** Fractional methylation change at CpG at the 4^th^ position within Cdx2 motif shows gain of methylation in *Eed^-/-^* cells at the sites with altered Cdx2 binding. **(E)** Heatmap showing chromatin accessibility dynamics at sites with gain or reduction of Cdx2 binding in *Eed*^-/-^ cells as compared to *WT* cells (as in a); chromatin accessibility from developmental timepoints at the Cdx2 bound loci is lost with lack of Cdx2 binding in adult cells (as seen in a) and rebinding of Cdx2 to such loci in *Eed^-/-^* cells reinduces open chromatin formation. **(F)** Genomic tracks showing recruitment of Cdx2 to developmental site in *Eed^-/-^* accompanied by gain of chromatin accessibility and increase in DNA methylation at the Cdx2 binding locus; dot plot at the bottom shows methylated (solid black dots) or unmethylated (white dots) status of the CpG in Cdx2 motif (4^th^ position) in 6 independent sequencing reads from *WT* and *Eed^-/-^*WGBS data. **(G)** Heatmap showing alterations in active enhancer mark H3K27ac at the sites with Cdx2 gain or reduction in *Eed*^-/-^ cells as compared to *WT* cells (as in a); enhancer activity from developmental timepoints at the Cdx2 bound loci is lost with lack of Cdx2 binding in adult cells (as seen in a) and rebinding of Cdx2 to such loci in *Eed^-/-^*cells reactivates these enhancers. **(H)** A model showing DNA methylation based Cdx2 binding at developmental target loci; Cdx2 is recruited to developmental sites upon gain of methylation of CpG (at 4^th^ position) within the Cdx2 motif in adult *Eed^-/-^* cells. See also Figure S5

To understand the effect of this flux in DNA methylation and Cdx2 binding on TF corecruitment, we conducted CUT&RUN for Ctcf. This showed that Cdx2 promoted Ctcf binding to developmental enhancers in *Eed^-/-^*cells (Figure 6A). Indeed, Ctcf signal at the loci with Cdx2 gains in *Eed^-/-^*cells reached almost the levels of its binding in native epithelial cells during developmental stages (Figure 6B). Moreover, 1,410 sites with significant gain and 511 with reduction of Ctcf binding in *Eed^-/-^* cells (*q* < 0.05, 1.5X) were associated with corresponding gain and reduction of Cdx2 binding (*p* < 0.001, Figure 6C, S6A-B). The sites where Ctcf is corecruited with Cdx2 in adult *Eed^-/-^* cells were highly linked to genes related to embryogenesis and pattern formation as compared to the loci that Ctcf binds without Cdx2, which showed association with genes involved in immune response and other homeostatic functions (Figure S6C). Together, these data strongly suggest that Cdx2 based priming and corecruitment controls Ctcf genomic binding and sustained presence at specific genomic sites in intestinal epithelial cells. On the other hand, while Hnf4a gets corecruited with Cdx2 at a few loci in *Eed^-/-^* cells, it’s binding shows some reduction in signal at the loci with reduced Cdx2 binding (Figure 6D), suggesting a role of Cdx2 in maintenance of Hnf4a binding in adult cells. To address the potential involvement of chromatin features other than the CpG methylation in the Cdx2 recruitment at developmental loci in adult *Eed^-/-^* cells, we looked at alterations to histone modifications in the *Eed^-/-^* cells. We see no significant change in H3K9me3, H2AK119ub, or H3K36me3, and a small number of loci (156 regions) with high H3K27me3 signal in the *WT* adult cells showed a gain of H3K36me2 signal in the *Eed^-/-^* cells (Figure 6E, S6D), suggesting that remethylation of CpG at the 4^th^ base position in the Cdx2 motif (Figure S2A) is a major driver of its recruitment to developmental target loci.

**Figure 6.**
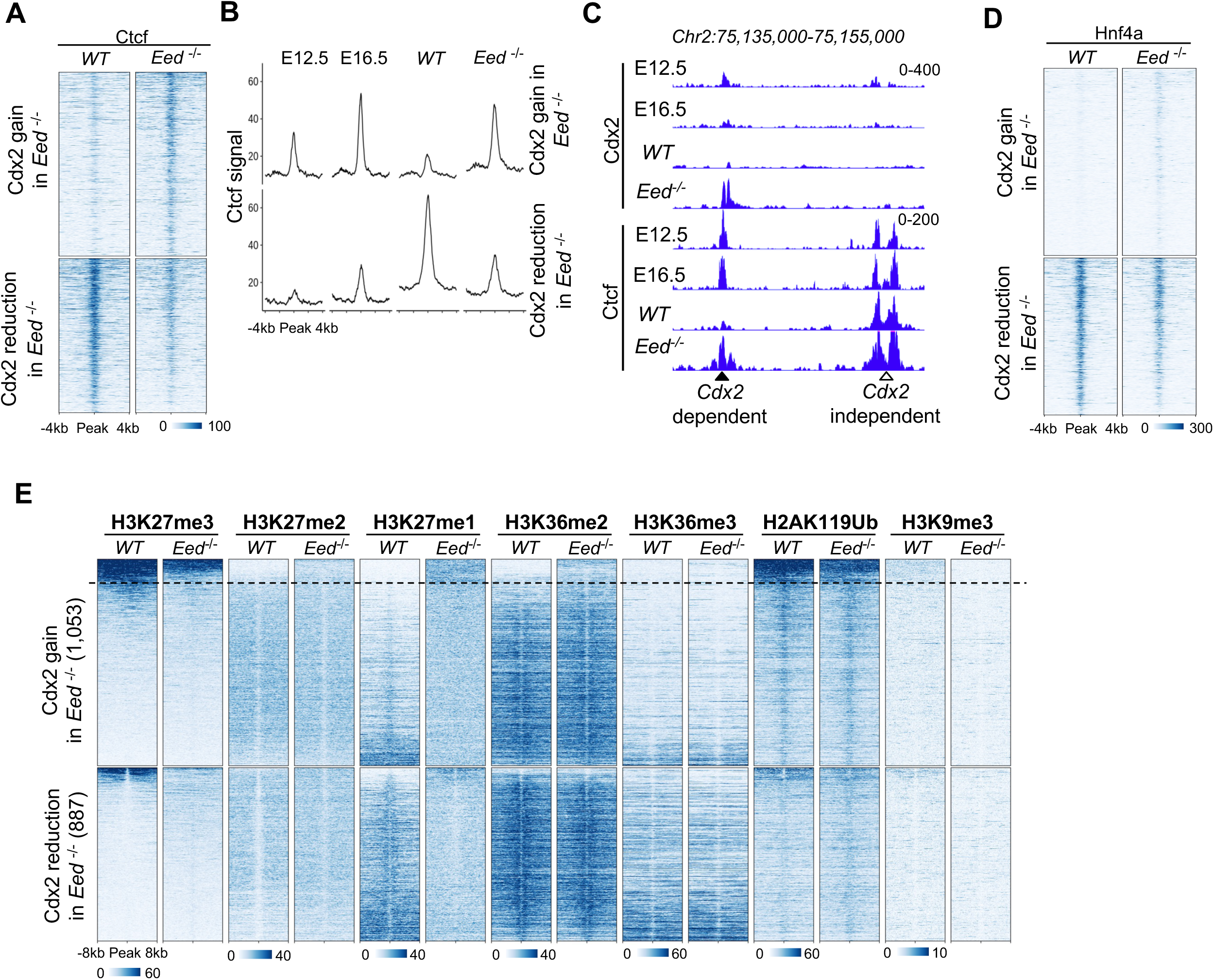
Perturbed Cdx2 binding resulting from DNA methylation changes leads to altered cobinding of TFs. **(A)** Heatmap showing Ctcf signal in adult villus and *Eed*^-/-^ epithelium at loci with gain or reduction of Cdx2 binding (as in Figure 5A). **(B)** Profile plots showing Ctcf signal from embryonic (E12.5), fetal (E16.5), adult villus, and *Eed*^-/-^ epithelium at the loci with gain or reduction of Cdx2 binding in *Eed*^-/-^ cells. **(C)** Genomic tracks showing corecruitment of Ctcf with Cdx2 in adult *Eed*^-/-^ cells reviving the early embryonic binding at E12.5; a nearby locus shows stable binding of Ctcf independent of Cdx2 occupancy. **(D)** Heatmap showing Hnf4a signal in adult villus and *Eed*^-/-^ epithelium at loci with gain or reduction of Cdx2 binding (as in Figure 5A). **(E)** Heatmaps showing signal of various histone modifications in adult epithelial cells at sites with Cdx2 gain or reduction in *Eed*^-/-^ cells (as in Figure 5A); regions in each cluster are arranged in decreasing order of the repressive H3K27me3 modification signal. The loci above the dotted line are positive for H3K27me3 and show no H3K36me2 signal in WT epithelium. See also Figure S6

### Biochemical modulation of DNA methylation causes altered Cdx2 binding

To further establish the effect of DNA methylation on Cdx2 binding, we altered CpG methylation more directly using a specific and well-established DNA methylation inhibitor GSK-3484862 (Azevedo Portilho et al., 2021) in HCT116 cells, a colorectal carcinoma cell line with a well- characterized epigenome including robust DNA methylation (Figure 7A) (Blattler et al., 2014). We first analyzed Cdx2 binding in duplicate experiments using CUT&RUN and observed Cdx2 occupancy at 10,197 loci (Macs2 peaks, *q* < 0.01), with the CpG-containing motif detected at 2,464 of these (FIMO, *p* < 0.005). Using triplicate WGBS data and a stringent cut-off of 5X coverage at the CpG in 4^th^ position of the Cdx2 motif, 471 sites displayed minimum 70% methylation at the Cdx2 bound motifs in untreated cells (Blattler et al., 2014). Exposure to 10µM GSK-3484862 for 6 days caused global loss of DNA methylation as detected by immunofluorescence for 5-methyl Cytosine (Figure 7B). CUT&RUN assays revealed a significant loss of Cdx2 binding at sites with the CpG-containing motif in treated cells (Figure 7C-D). Importantly, removal of the inhibitor from culture media for 3 days, which allows recovery of the DNA methylation (Butz et al., 2022), caused significant rebinding of Cdx2 demonstrating a DNA methylation based recruitment of Cdx2 (Figure 7C-E); 500 randomly chosen Cdx2 peaks without a CpG within the detectable motifs showed no significant alteration in Cdx2 binding upon the inhibitor treatment or upon removal of the inhibitor (Figure 7F). These results demonstrate that the recruitment and the dynamics of Cdx2 binding is controlled by its motif distribution and alterations of CpG methylation.

**Figure 7.**
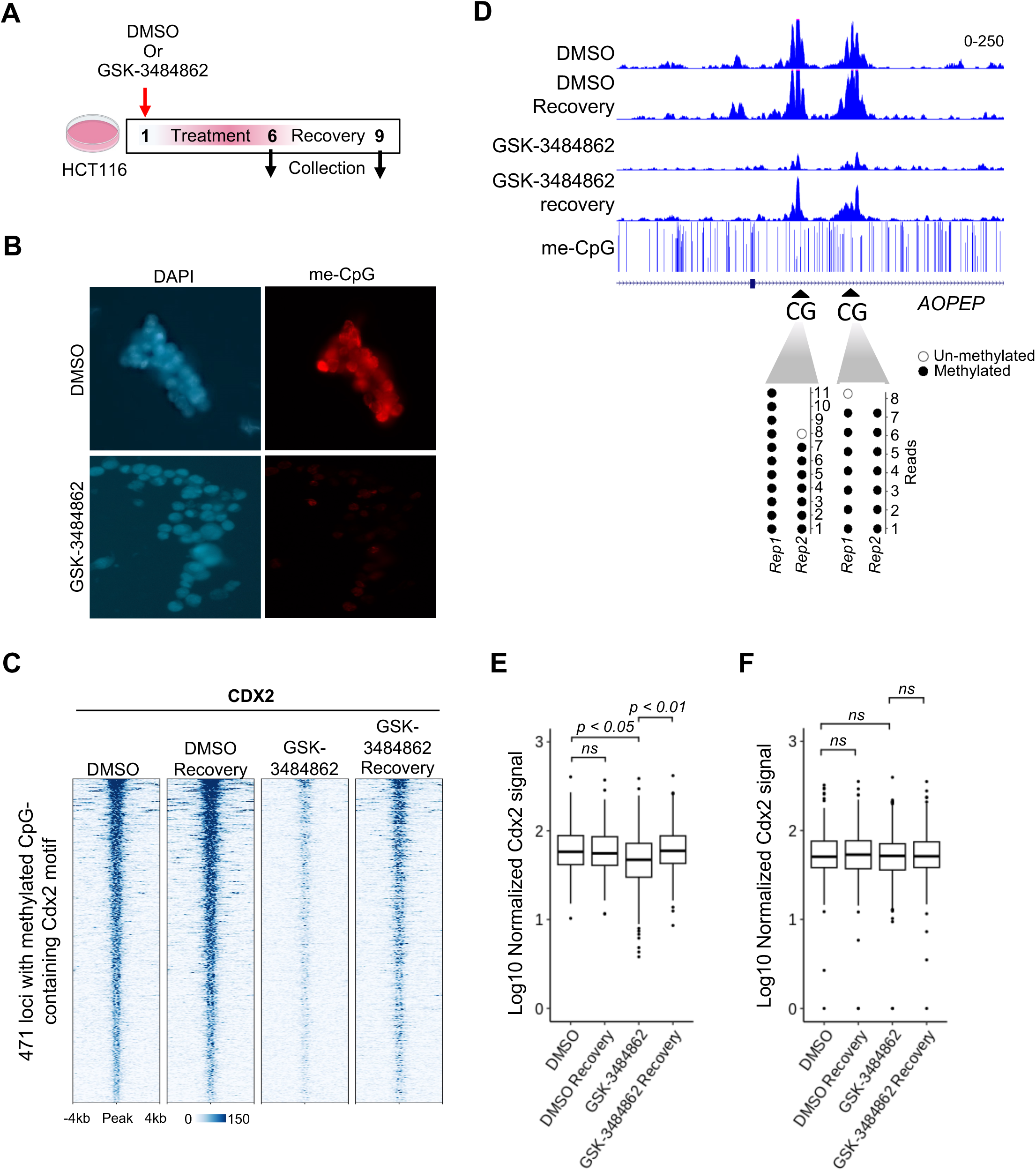
Direct inhibition of DNA methylation causes loss of Cdx2 binding and remethylation at binding loci allows its recruitment. **(A)** Experimental schematic showing treatment of HCT116 cells with DNA methylation inhibitor GSK-3484862 or DMSO (control) for 6 days in order to eliminate CpG methylation followed by 3 days of recovery without the drug (or DMSO) to allow for remethylation. **(B)** Fluorescence micrographs showing loss of DNA methylation upon treatment with inhibitor GSK-3484862 for 6 days. **(C)** Heatmap showing loss of Cdx2 binding at loci with CpG-containing Cdx2 motif in HCT116 cells treated with DNA methylation inhibitor as compared to DMSO (control); upon removal of the inhibitor, Cdx2 is recruited back to these loci within 3 days. **(D)** Representative genomic tracks showing loss of Cdx2 binding upon elimination of DNA methylation in HCT116 cells through treatment with GSK- 3484862 as opposed to the stable binding upon treatment with DMSO (control); removing the inhibitor from the cell culture media causes recovery of the Cdx2 binding. **(E)** Box and whisker plot showing significant loss of Cdx2 binding at 471 loci with CpG-containing Cdx2 binding motif upon treatment with DNA methylation inhibitor and recovery of this binding signal upon removal of the inhibitor; there is no significant change in Cdx2 binding at 500 randomly chosen loci without the CpG-containing motif **(F)**.

## DISCUSSION

### Distinct mechanisms of pioneer TF action in developing and adult tissues

Pioneer TFs establish tissue patterning and function through the establishment of tissue-specific gene expression networks. To do this, they rapidly activate other TFs, which in turn allows expansion of the lineage and cell type-specific enhancer landscape over time (Zaret, 2020). Our data show that Cdx2 activates a proportionally high number of TF genes such as *Sox4* during early development (Figure 1D), and it binds directly to promoters of such TF genes and other developmental targets. By activating other TFs through simple promoter-based control during development, pioneer factors may rapidly increase the potential to establish lineage-specific enhancers necessary for more complex gene control required by adult homeostatic tissue functions. Indeed, recent studies looking at TF action in developmental and adult erythroblast cells show that simple promoter-centric regulation through Gata1 targets embryonic-specific genes, while increasing combinatorial enhancer-driven gene control through Myb in adult cells (Cai et al., 2020). We show that such promoter-based targeting of Cdx2 during intestinal development may be programmed in elevated presence of the CpG containing motif at the developmental target sites in comparison to the canonical motif without CpG. Thus, for pioneer TFs like Cdx2 with variable affinity for different motifs, the relative abundance of such motifs may be a potential way to shift from rapid promoter-based regulation in development to a more nuanced enhancer-driven control in adult tissue. Interestingly, the Cdx2 loci specifically bound in development show more conservation across species as compared to the loci bound only in adult cells. Together, these data suggest a conserved molecular mechanism based on DNA methylation for TF recruitment to developmental targets.

### DNA methylation sensitivity and control of TF cobinding

Along with DNA sequence, CpG methylation and chromatin structure can affect TF binding (Crispatzu et al., 2021; Isbel et al., 2022; Nguyen et al., 2016). Ctcf has a relatively large motif with multiple CpGs, which makes it sensitive to DNA methylation (Holwerda and de Laat, 2013; Maurano et al., 2015). A small proportion of all known Ctcf binding sites are highly conserved across species and show high occupancy of Ctcf across tissues, while the tissue specific Ctcf binding sites show low occupancy (Essien et al., 2009). Although DNA methylation and chromatin remodelers such as ISWI are involved in Ctcf recruitment (Wang et al., 2012a; Wiechens et al., 2016), the role of cobinding TFs, which may allow Ctcf to overcome the DNA methylation barrier and facilitate its tissue-specific binding, remains understudied. Interestingly, the Ctcf motif is not highly detectable in the developmental clusters Dev 1-3 (Figure 2A). In this light, cobinding of Ctcf with Cdx2 in the developing intestinal epithelium (Figure 2E) suggests a developmental stage-specific mechanism where the pioneer TF Cdx2 and its chromatin sensitivity allows Ctcf occupancy through corecruitment. This Cdx2 based recruitment of Ctcf is further highlighted in adult cells when gain of DNA methylation at developmental Cdx2 binding loci results in ectopic recruitment of Cdx2 and cobinding of Ctcf. On the other hand, the loci occupied by Cdx2 specifically in adult cells (Adult 1-3) show strong presence of an Hnf4a motif and co-binding of Hnf4a, and reduction of Cdx2 at these loci reduces Hnf4a occupancy. These results suggest that Cdx2, through its motif variety and distribution as well as chromatin sensitivities, may control access and maintenance of Ctcf and Hnf4a access at distinct target sites.

### Motif variability, distribution, and DNA methylation control TF recruitment and sustained binding

Although *in vitro* studies suggest that many TFs have DNA methylation sensitivity, how DNA methylation changes during tissue development direct TFs to their cell type and developmental stage specific target sites is not well understood. In the *in vivo* context, both DNA methylation and TF binding may be influenced by nucleosome distribution, histone modifications, motif variability and frequency, and other cobinding TFs. Moreover, these factors may have a distinct impact on TF recruitment versus their sustained binding. Many TFs that prefer DNA methylation mediated recruitment seem to mediate demethylation at their target loci upon binding (Stadler et al., 2011). This is also true for Cdx2, as it binds its developmental targets in intestinal epithelium (E12.5 and E16) when the CpG in Cdx2 motif is uniquely methylated, which then gets demethylated in adult cells. This suggests that Cdx2 may recruit demethylases at these loci through direct or indirect interactions. Indeed, Ctcf, which shows cobinding with Cdx2 at many such loci (Figure 3E) can inhibit DNA methylation by locally suppressing the activity of the ubiquitously expressed DNA (cytosine-5)-methyltransferase 1 (DNMT1) through cobinding with poly(ADP-ribose) polymerase 1 (PARP1), an enzyme that adds ADP–ribose groups to DNMT1 to inactivate it (Guastafierro et al., 2008; Stadler et al., 2011; Zampieri et al., 2012).

In multiple studies, we have previously shown that embryonic and fetal enhancers remain hypomethylated in adult cells and they are ectopically activated upon loss of PRC2 action and the repressive H3K27me3 modification (Jadhav et al., 2019; Jadhav et al., 2020). In the same mouse model system, we now find that induced loss of PRC2 activity causes remethylation of CpG within the Cdx2 motif and induces Cdx2 recruitment to developmental enhancers in adult cells. In this light, our results reveal a dual-purpose molecular mechanism where CpG methylation facilitates Cdx2 recruitment at developmental loci, while subsequent demethylation at these sites precludes Cdx2 binding in adult cells.

Overall, our data unveil that through the capability to bind motifs with and without CpGs distributed unequally at developmental and adult target sites, respectively, Cdx2 is able to navigate distinct DNA methylation profiles and recruit other TFs to establish lineage specific enhancer patterns.

## METHODS

### EXPERIMENTAL MODEL AND SUBJECT DETAILS

*ROSA26^LSL-tdTomato^* mice were purchased from The Jackson Laboratories; *Eed^fl/fl^* mice (Xie et al., 2014) and *Villin^CreER-T2^*(el Marjou et al., 2004) were generous gifts from S. Orkin (Boston Children’s Hospital) and S. Robine (Institut Pasteur, France), respectively. Mice were handled and treated according to protocols approved by the Department of Animal Resources at the University of Southern California. Mice were housed in a facility maintained at 23 ±1°C, 50 ±10% humidity, and 12 hr light/dark cycles. Genetically altered alleles and were confirmed by genotyping mice using PCR at weaning and during experiments.

### Mouse treatments

For induced deletion of *Eed*, mice with age 8 weeks or older were given 2 mg tamoxifen using intraperitoneal (i.p.) injections on 5 consecutive days, and tissues were harvested at times as indicated for individual experiments.

### Isolation of intestinal epithelial cells

Adult villus cells were procured using proximal 1/3^rd^ small intestine. Immediately after harvesting, the tissue was cut along the length to expose epithelium and washed with cold phosphate-buffered saline (PBS). Tissue was rotated in 5 mM EDTA in PBS (pH 8) at 4°C for 30 min, with vigorous manual shaking every 10 min. The recovered epithelium was filtered using 70 *µm* filters and the villi retained on the filter were collected in ice-cold PBS. Villi were collected by centrifugation at 300 *g* for 5 min and single-cell suspension was generated by rotating in 20ml of 4% TrypLE solution (Invitrogen) in DMEM at 37°C for 30 min; 15ml DMEM was added to neutralise the TrypLE solution and cells were collected using centrifugation. For the isolation of epithelial cells at prenatal timepoints E12.5 and E16.5, embryos were harvested from pregnant mice 12 and 16 days after identification of copulation plug, respectively. The small intestine (between stomach and cecum) was chopped into small pieces with razor and digested in 500ul of 0.25% trypsin for 20 minutes at 37 °C to generate single cell suspension. Reaction was stopped by addition of 500ul of DMEM with 2% FBS. Cells were spun down at 500 *g* for 5 minutes at 4 °C and resuspended in PBS with 2% BSA. EpCam+ epithelail cells were isolated by EasySep™ Release Mouse Biotin Positive Selection Kit according to manufacturer protocol and EpCAM ab (Invitrogen 13-5791-82).

### Native-ChIP-seq

Native-ChIP-seq was performed as previously described (Lorzadeh et al., 2016; Lorzadeh et al., 2017). In brief, cells were lysed in 0.1 % Triton X-100 and 0.1% Sodium Deoxycholate with protease inhibitor cocktail. Chromatin in the cell lysate was digested using Microccocal nuclease (MNase, New England BioLabs, M0247S) at room temperature for 5 min and 5 ul 0.25 mM EDTA was added to stop the reaction. The digested chromatin were incubated with anti-IgA magnetic beads (Dynabeads,Themo Fisher, 10001D) for 2 h for pre-clearing and then incubated overnight with antibody-bead complexes with antibodies against H3K27me3, H3K27me2, H3K27me1, H3K36me2, H3K36me3, and H2AK119Ub (Diagenode, C15410195, C15410046, C15410045, C15310127, C15410192 8240S, respectively) in immunoprecipitation (IP) buffer (20 mM Tris-HCl pH 7.5, 2 mM EDTA, 150 mM NaCl, 0.1% Triton X-100, 0.1 % Sodium Deoxycholate) at 4 °C. IPs were washed 2 times by Low Salt (20 mM Tris-HCl pH 8.0, 2 mM EDTA, 150 mM NaCl, 1% Triton X-100, 0.1% SDS) and High Salt (20 mM Tris-HCl pH 8.0, 2 mM EDTA, 500 mM NaCl, 1% Triton X-100, 0.1% SDS) wash buffers. IPs were eluted in elution buffer (1% SDS, 100 mM Sodium Bicarbonate) for 1.5 h at 65 °C. Histones were digested by Protease (Qiagen 19155) for 30 min at 50 °C and DNA fragments were purified using Sera Mag magnetic beads in 30% PEG. Illumina sequencing libraries were generated as previously described (Lorzadeh et al., 2022) by end repair, 3′ A-addition, and Illumina sequencing adaptor ligation (New England BioLabs, E6000B- 10). Libraries were then indexed and PCR amplified (10 cycles) and sequenced on Illumina HiSeq 2500 sequencing platform following the manufacture’s protocols (Illumina, Hayward CA).

### Whole genome bisulfite sequencing (WGBS)

MasterPure DNA purification kit (Epicenter MCD85201) was used to purify genomic DNA from the cells followed by bisulfite conversion using 50 ng DNA along with the EZ DNA Methylation-Gold kit (Zymo Research D5005). 10 ng of bisulfite-converted DNA was used to generate the whole genome bisulfite sequencing (WGBS) libraries with the EpiGenome Methyl-Seq kit (Epicenter EGMK81312). We used AMPure beads (Beckman Coulter) to purify the library material and confirmed that the library DNA was in the size range of 200-800 bp using high sensitivity DNA Chip detection (Bioanalyzer 2100, Agilent Genomics), and sequenced the libraries on a NextSeq 500 instrument (Illumina) to generate 150-bp paired-end reads; up to 50% PhiX phage DNA (Illumina) was mixed with the libraries to assist the sequencing.

### CUT&RUN

CUT&RUN was performed using EpiCypher CUTANA kit (14-1048) according to manufacturer’s protocol with the following changes. Freshly isolated epithelial cells were resuspended in the permeabilization buffer (CUTANA Wash Buffer, 0.01% Digitonin, and 1X protease inhibitor) and incubated on ice for 5 minutes and captured on Concanavalin A beads at room temperature for 10 minutes. Captured cells were resuspended in 180ul of antibody buffer (permeabilization buffer with 2mM EDTA) with 0.5 ug of antibody against Cdx2 (Cell Signalling, D11D10), Ctcf (Diagenode, C15410210), or Hnf4a (Santa Cruz Biotechnology, sc-374229) and incubated overnight. Antibody bound chromatin was digested at 4 °C for 30 minutes on a nutator. Fragment release postdigestion was done at 4 °C for 30 minutes and purified using SeraMag beads with 30% PEG-8000. Library construction was done as described in the Native-ChIP-seq section.

### ATAC-seq

50,000 *Eed^-/-^* adult villus cells were lysed in 50 µl cold ATAC lysis buffer (10 mM Tris·Cl, pH 7.4, 10 mM NaCl, 3 mM MgCl2, 0.1% (v/v) Igepal CA-630) followed by centrifugation at 500 *g* at 4°C to isolate nuclei. Nuclei were resuspended in 50 µl transposition reaction mix with Nextera Tn5 Transposase (Illumina, FC-121-1030) and incubated for 30 min at 37°C. Transposed DNA was purified using columns (Qiagen, 28004) and amplified using high-fidelity 2X PCR Master Mix (New England Biolabs) using primers with standard ATAC-seq barcodes (Buenrostro et al., 2015). AMPure beads (Beckman Coulter, A63880 were used to remove primer dimers and libraries were sequenced on a NextSeq 500 instrument (Illumina) to generate 75 bp single-end reads.

### Cross-linked ChIP-seq

∼1×10^6^ wild type and *Eed^-/-^* villus cells were fixed in 1% formaldehyde for 25 min at room temperature immediately after isolation. The reaction was stopped by addition of Glycine to a final concentration of 0.13 M. Cells were lysed using a buffer containing 30 mM Tris-HCl (pH 8), 1% SDS, 10 mM EDTA, and protease inhibitors (Roche), and the chromatin was sonicated for 50 min using a Covaris sonicator (5-min on/off cycles at 4 °C). Debris were removed using centrifugation and the chromatin was incubated overnight at 4°C with antibody against H3K9me3 (Abcam, Ab8898). Antibody-bound chromatin was captured using magnetic beads (Themo Fisher, 10001D) and washed using low-salt (20 mM Tris-HCl pH 8.1, 150 mM NaCl, 2 mM EDTA, 0.1% SDS, 1% TritonX-100), high-salt (20 mM Tris-HCl pH 8.1, 500 mM NaCl, 2 mM EDTA, 0.1% SDS, 1% TritonX-100), and lithium chloride (10 mM Tris pH 8.1, 0.25 M LiCl, 1 mM EDTA, 1% NP-40, 1% deoxycholate) buffers. Chromatin was treated with 1% SDS and 0.1 M NaHCO_3_ for 6 h at 65°C to reverse the cross-links and DNA was purified using columns (Qiagen) followed by ChIP-seq library preparation using ThruPLEX kit (Rubicon, R400427) and sequenced using NextSeq 500 instrument (Illumina) to produce 75-bp single-end reads.

### QUANTIFICATION AND STATISTICAL ANALYSIS

### Computational analyses

#### RNAseq

Raw RNA-seq reads were aligned to the mouse genome (GRCm38, GENCODEv25) using STAR aligner v2.7.8a (Dobin et al., 2013) in two-passed mode followed by RSeQC v2.6.2 assessment (Wang et al., 2012b) to determine per-base sequence quality, per-read GC content (∼50%), comparable read alignments to +/- strands, exon vs intron read distributions, and 3’ bias. Transcript levels were expressed as read counts using HTSeq v0.13.5 (Anders et al., 2015) and normalized across libraries using DESeq2 (Love et al., 2014), followed by conversion into reads per kb of transcript length per 1M mapped reads (RPKM). We determined differential expression between samples using DESeq2, with false-discovery rate (FDR) as indicated in the text. All alignments and initial processing steps were run using the Snakemake workflow management system (v6.6.1). Cartoon Illustrations were created with BioRender.com.

#### ChIP-seq and ATAC-seq

ChIP-seq and ATAC-seq data were trimmed using trim-galore (v0.6.6) to remove adapters and low-quality reads. Trimmed sequences were aligned using bwa-mem (v0.7.17) with default parameters. Non-uniquely mapped reads, duplicate reads, and reads with a MAPQ score of less than 5 were removed using Samtools (v1.17). Peak calling was done using MACS (v2.2.7.1) with a q-value cutoff of 0.01 in sharp mode for H3K27ac and ATAC, and broad mode for H3K27me3,2 and 1, H3K36me3 and 2, H2AK119Ub and H3K9me3. BigWig files were generated using deeptools (v3.5.1) with a bin size of 10, smooth length of 30, and quantile normalized using HayStack. Bedtools (v2.30.0) was used to removed blacklisted regions defined by ENCODE blacklist.

#### Analysis of DNA methylation data

Whole genome bisulfite data were aligned to mm10 in non-strand specific mode using Bismark v0.23.0 (Krueger and Andrews, 2011). Coverage from both strands for each CpG were combined and fractional methylation was calculated for each CpG using custom AWK command. Differential single CpGs were identified using fisher exact test (p < 0.001). Differential CpG with at least 20% difference in fractional methylations and within 300bp of one another were merge using custom AWK command. Hypermethylated regions with less than 40% methylation were discarded.

#### Motif enrichment analysis

Motif enrichment comparison between developmental and adult Cdx2 sites were conducted using HOMER findMotifGenome with default parameters across the whole Cdx2 binding site. Instance of the motif were identified as previously described (Lorzadeh et al., 2022). In brief, significant instances of the Cdx2 motif (*p <* 0.005) was identified using MEME-SUIT fimo command (Grant et al., 2011). Instances of CpG containing Cdx2 motif were further filtered to include motif with CpG at the 4^th^ position and CGTA sequence at 4 to 7^th^ positions. Both 10bp and 12bp CpG containing Cdx2 motif (Figure S2A) were considered together for counting of CpG containing motifs.

#### Identification of unmethylated (UMR) and low-methylated (LMR) regions

MethylSeekR package (Burger et al., 2013) was used along with the mouse bisulfite-converted genome (BSgenome.Mmusculus.UCSC.mm10) from Bioconductor (www.bioconductor.org) as the reference to determine UMRs and LMRs. We only used CpGs with coverage_5 or more and eliminated C nucleotides overlapping known SNPs between the reference strains (C57BL/6J and 129/S5) to avoid any effects of polymorphism. Methylation levels were smoothed over 5 consecutive CpG dinucleotides hypomethylated regions were identified as UMRs (unmethylated and minimum 5 CpGs and less than 10% fractional methylation) or LMRs (minimum 5 CpGs with fractional methylation between 10% and 60%).

## CONTACT FOR REAGENT AND RESOURCE SHARING

Requests should be directed to the Lead Contact, Unmesh Jadhav. Requests will be fulfilled by after execution of a suitable Materials Transfer Agreement.

## DATA AND SOFTWARE AVAILABILITY

The sequencing data reported in this study is accessible through Gene Expression Omnibus (GEO). The accession number for the data reported is GSE253736.

## ACKNOWLEDGEMENTS

This work was supported by NIH grants K01DK113067 and R03DK134799 to U.J. A.L was supported by Canadian Institute of Health Research (CIHR) Post-Doctoral Fellowship.

## CONTRIBUTIONS

A.L. and U.J. conceptualized the study design. A.L. and G.E. performed the analysis. A.L. and S.S. performed experimental work. A.L. and U.J. wrote the manuscript. All authors read and approved the final manuscript.

---

**Figure S1, Related to Figure 1.**
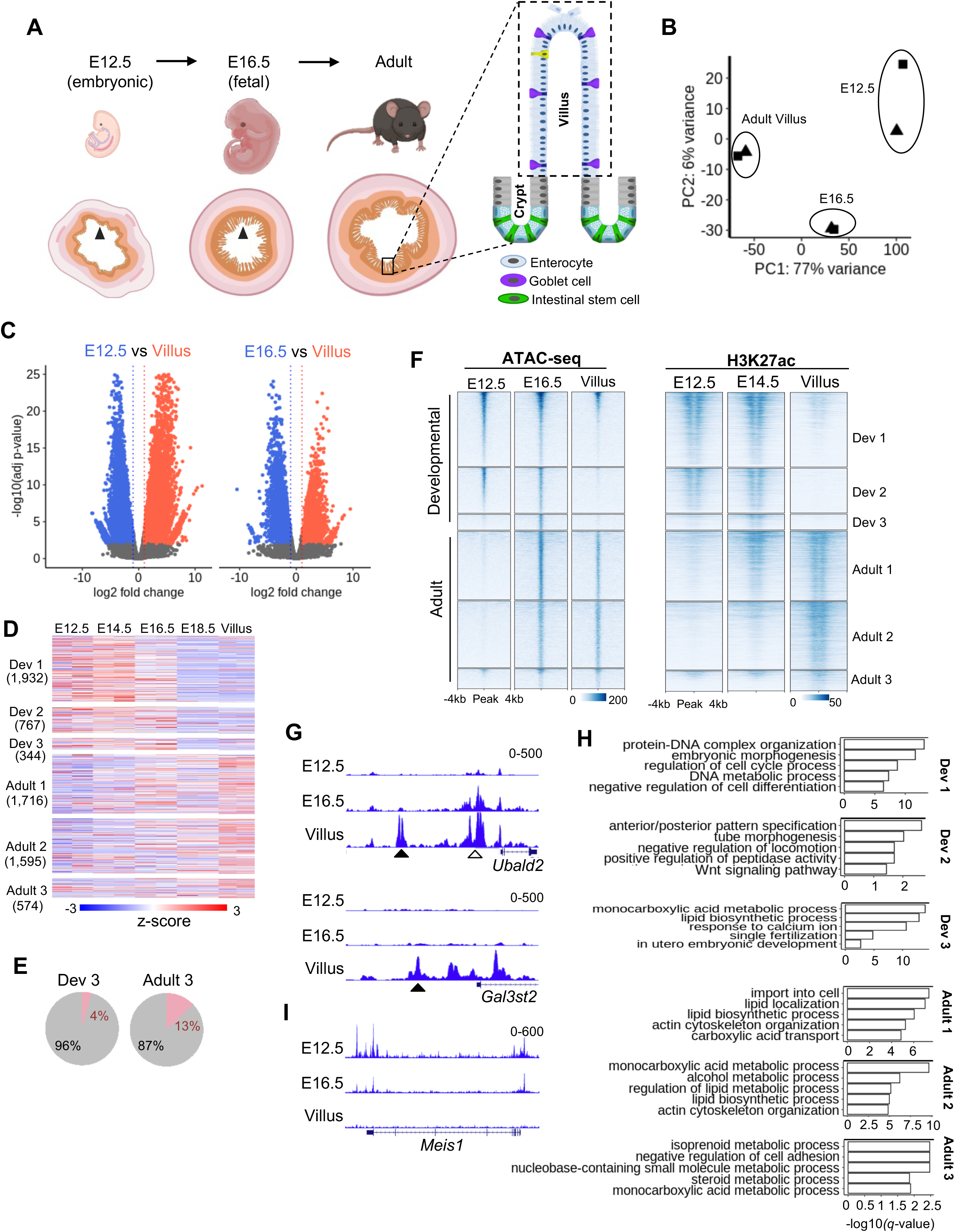
**(A)**Drawings depicting different stages of mouse intestinal development; epithelial cells lining the intestinal lumen (arrowheads) and adult post-mitotic villus cells (predominantly enterocytes) were purified to profile Cdx2 binding. **(B)** Principal component analysis showing high concordance among replicates of Cdx2 CUT&RUN data for embryonic (E12.5), fetal (E16.5), and adult villus epithelial cells. **(C)** Volcano plots showing significant (*q* < 0.01, 2-fold change) gains (red) or losses (blue) of Cdx2 binding in villus as compared to E12.5 (left panel) and E16.5 (right panel) epithelial cells. **(D)** Heatmap representing expression of genes near (+/- 25 kb) dynamic Cdx2 binding sites (Dev 1-2 and Adult 1-3 in Figure 1A) in developing epithelial cells E12.5, E14.5, E16.5, E18.5, and adult villus; numbers in bracket represent genes found near loci in the respective subgroups. **(E)** Pie charts showing percentages of Cdx2 binding sites in Dev 3 and Adult 3 subgroups shown in Figure 1A within and outside promoters (TSS -2 kb to +1 kb). **(F)** Heatmap showing chromatin accessibility (ATAC-seq) (Jadhav et al., 2019) and active promoter and enhancer associated H3K27ac signal (Kazakevych et al., 2017) at Cdx2 bound loci (as in Figure 1A) during developmental timepoints and in adult villus. **(G),** Genome browser tracks showing recruitment of Cdx2 at *Ubald2* and *Gal3st2* genes beginning at E16.5 (white arrowhead) or in villus (black arrowhead) cells, respectively. **(H),** Pathway enrichment analysis of genes associated with dynamic Cdx2 binding sites (+/- 25 kb from Dev 1-3 and Adult 1-3 in Figure 1A) at E12.5, E16.5, and adult villus epithelial cells. **(I)** Genome browser view showing Cdx2 binding at *Meis1* gene in E12.5, E16.5, which is lost in adult villus cells.

**Figure S2. Related to Figure 2.**
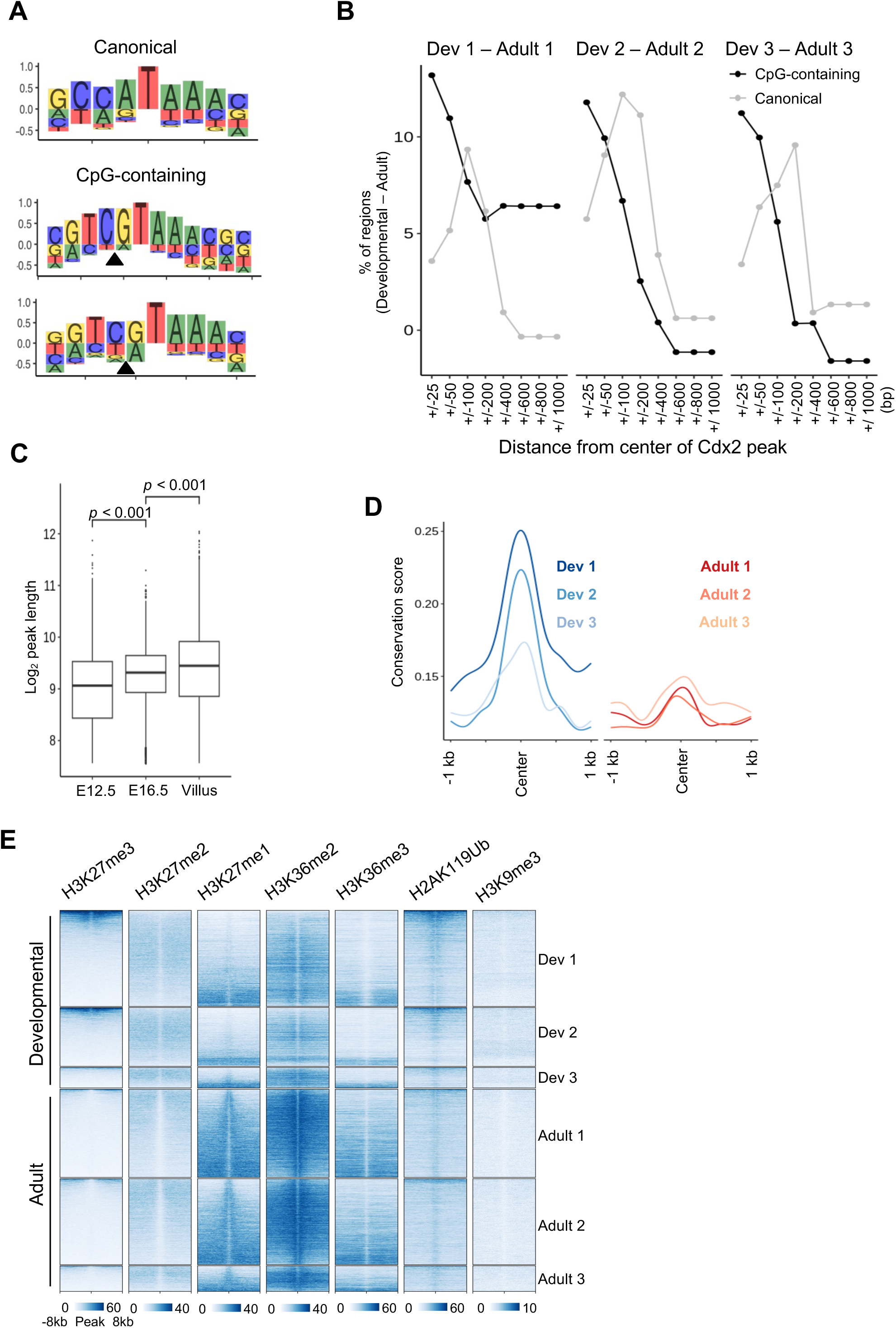
**(A)** Canonical (non-CpG-containing) and CpG-containing Cdx2 motifs; black arrowheads indicate CpG located at the 4^th^ base position within the motif. **(B)** Plots showing that comparatively larger proportion of regions with embryonic and fetal binding of Cdx2 (Dev 1-3) have presence of CpG-containing motif as compared to regions with adult cell specific binding (Adult 1-3); the CpG-containing motif is more prevalent near the center of Cdx2 binding peak (+/- 50 bp), while the canonical motif is more readily detected beyond 50 bp form the peak center. **(C)** Box plot showing differences in average width of the Cdx2 peak in E12.5, E16.5, and villus epithelial cells. **(D)** Profile plots showing cross-species conservation of various genomic loci with developing or adult epithelium specific (Dev 1-3 and Adult 1-3, respectively) Cdx2 binding. **(E)** Heatmaps showing signal of various histone modifications in adult epithelial cells at sites with dynamic Cdx2 binding through epithelial maturation (as in Figure 1A); the regions in each cluster are arranged in decreasing order of the repressive H3K27me3 modification signal.

**Figure S3. Related to Figure 3.**
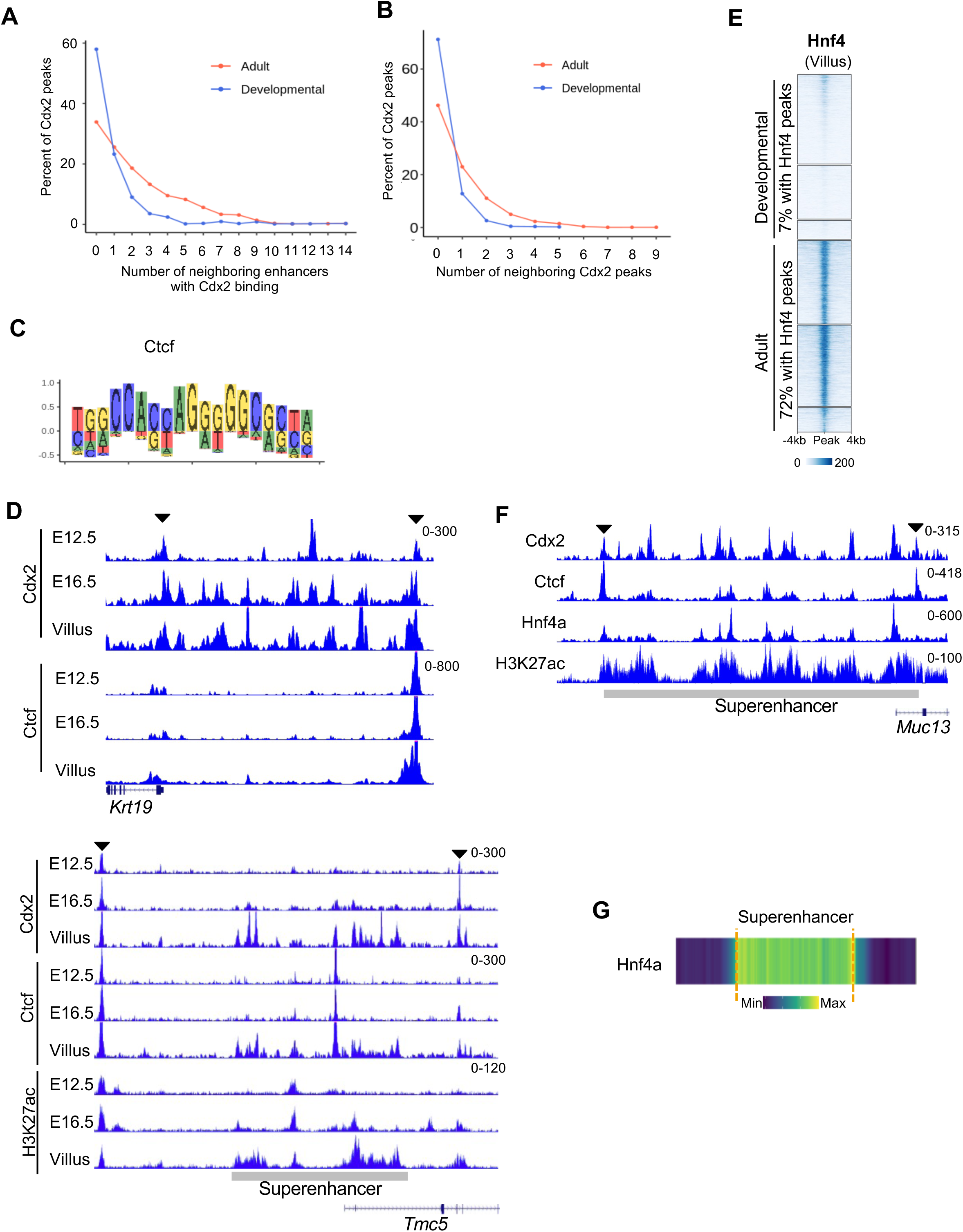
**(A)** Percentage of developmental (blue) or adult (red) cell specific Cdx2 bound enhancers with designated number of other Cdx2 bound enhancers nearby (+/- 50 kb). **(B)** Percentage of developmental (blue) or adult (red) cell specific Cdx2 bound loci (promoter or enhancer) with designated number of other Cdx2 peaks within 10 kb distance upstream or downstream. **(C)** Ctcf motif showing multiple CpGs. **(D)**, Genome browser view of Cdx2 and Ctcf signals near genes *Krt19* and *Tmc5* showing Cdx2 and Ctcf binding defining boundaries (or region, indicated by black arrowheads); H3K27ac signal near *Tmc5* shows superenhancer activity in adult cells that grows with the Cdx2 and Ctcf binding in fetal (E16.5) development. **(E)** Heatmap showing Hnf4a signal in adult villus cells at developmental and adult cell specific Cdx2 binding sites; the regions are displayed in decreasing order of Cdx2 signal as in Figure 1A. 7% and 72% loci with development and adult cell specific Cdx2 binding, respectively, show Hnf4a binding. **(F)** Genome browser view of Cdx2, Ctcf, Hnf4a, and H3K27ac signals at an adult superenhancer showing spread of Cdx2 and Hnf4a binding within the boundaries of the superenhancer. **(G),** Heatmap showing Hnf4a signal at superenhancers found in adult villus. Superenhancer boundaries are shown with a dotted orange line.

**Figure S4. Related to Figure 4.**
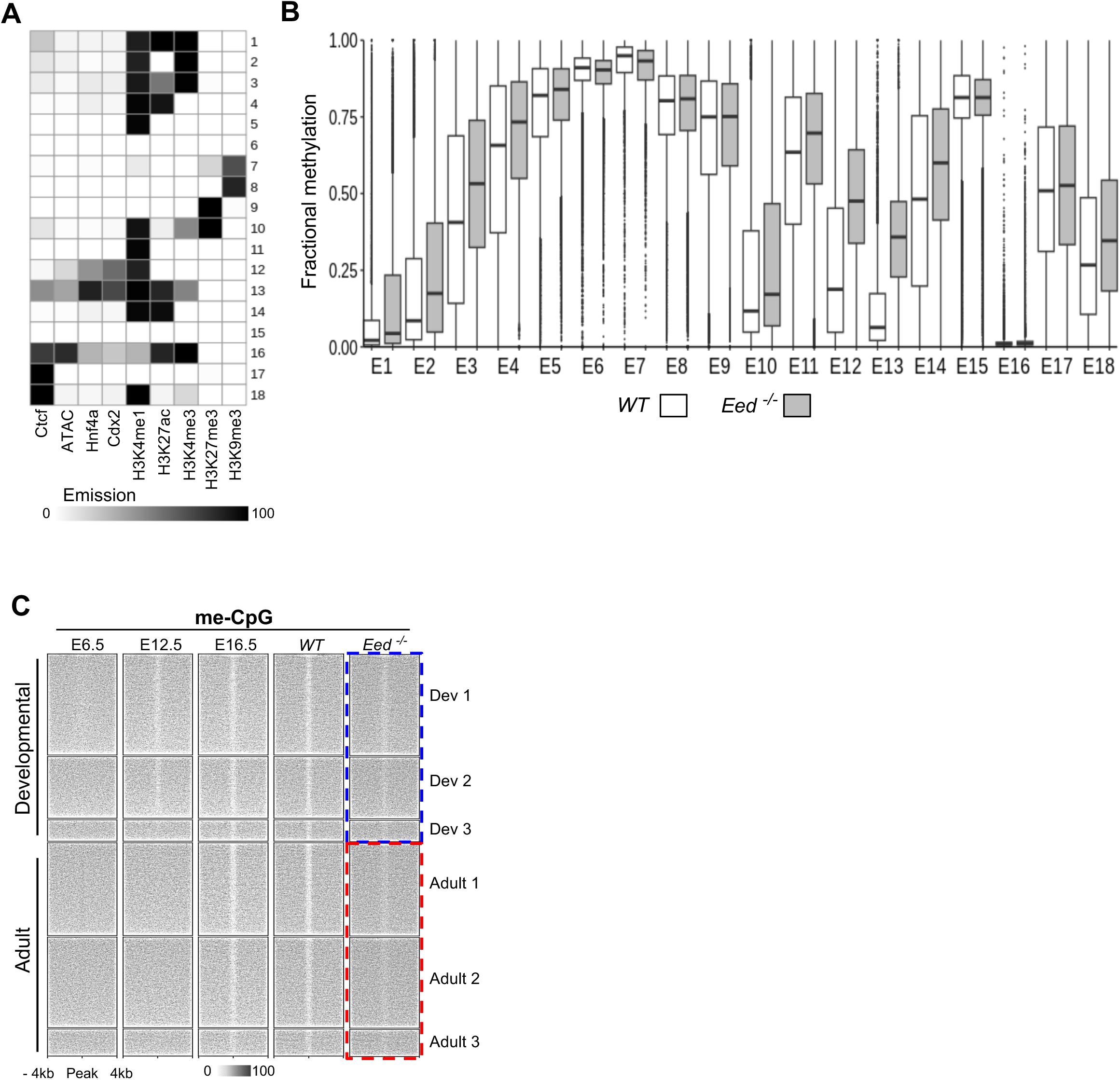
**(A)** Heatmap showing 18 different genomic states identified by CHROMHMM based on TF binding and histone modifications. **(B)** DNA methylation changes at these loci upon loss of PRC2 activity (*Eed^-/-^*). **(C)** Heatmap showing DNA methylation at Cdx2 binding loci in early endoderm (E6.5) (Seisenberger et al., 2012), and embryonic (E12.5), fetal (E16.5), and adult epithelial (villus) cells (as in Figure 2F) (Jadhav et al., 2019); induced loss of *Eed* and PRC2 action causes increased methylation at developmental (Dev 1-3) and adult (Adult 1-3) hypomethylated loci (blue and red dotted boxes, respectively).

**Figure S5. Related to Figure 5.**
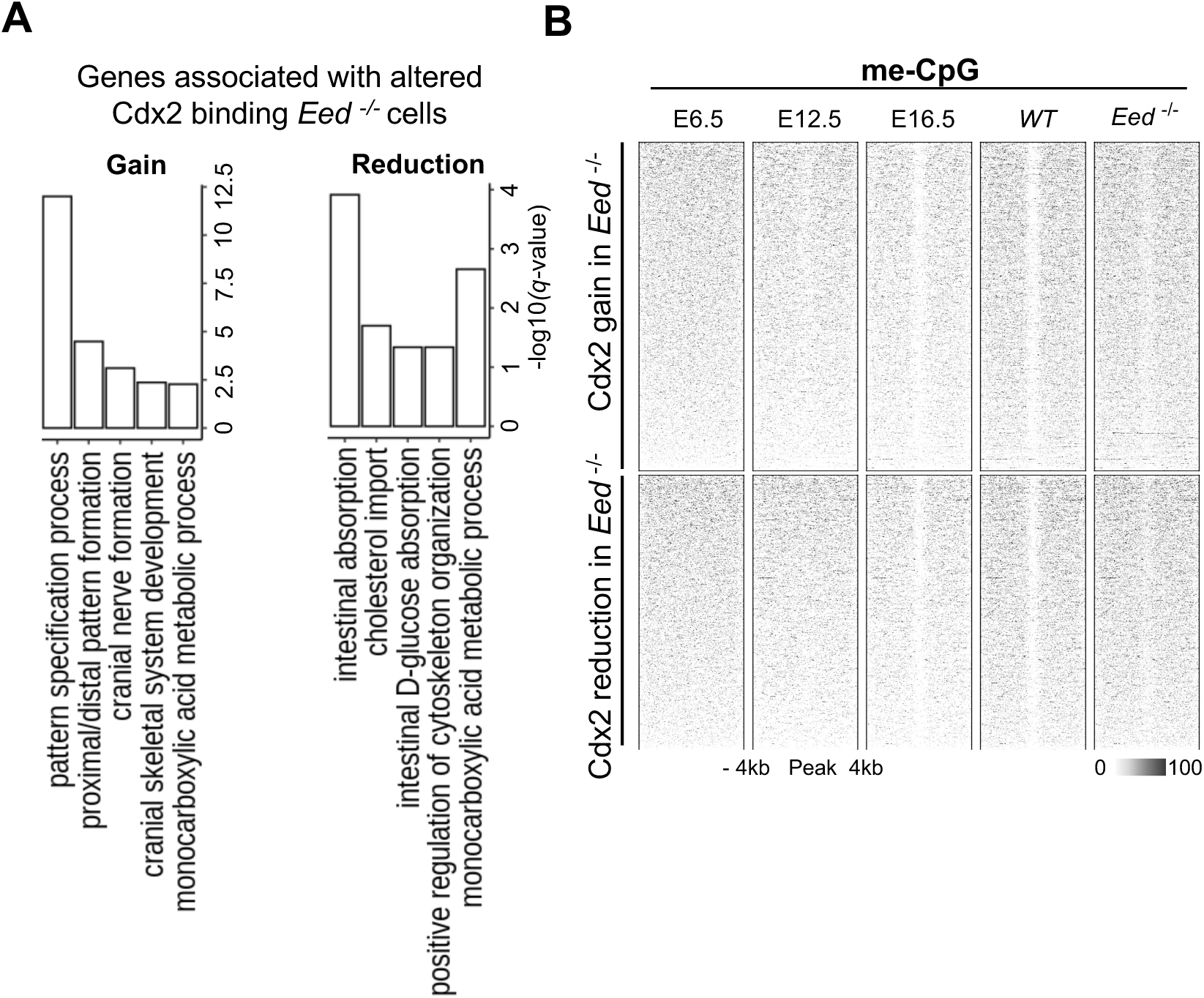
**(A)** Gene ontology analysis for genes within 50 kb of the loci with altered Cdx2 binding in *Eed^-/-^* cells highlights the recruitment of Cdx2 to developmental targets upon loss of PRC2 activity. **(B)** Heatmap showing DNA methylation dynamics at sites with gain or reduction of Cdx2 binding in *Eed*^-/-^ cells as compared to *WT* cells (as in Figure 5A).

**Figure S6. Related to Figure 6.**
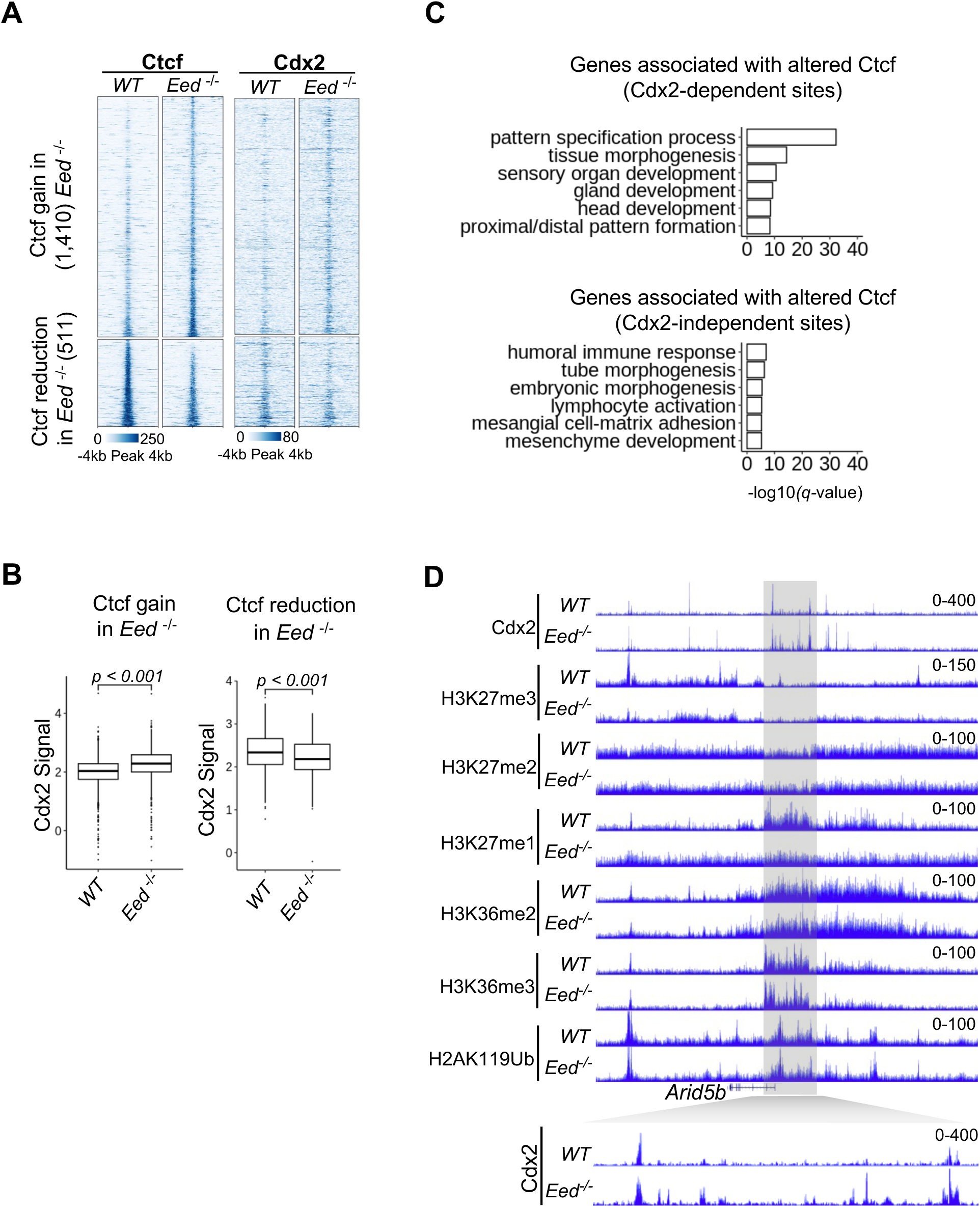
**(A)** Heatmap showing significant (*q* < 0.05, 1.5×) gain and reduction of Ctcf binding in *Eed*^-/-^ cells as compared to the native villus cells; corresponding heatmap of Cdx2 signal at these loci shows corecruitment of the two TFs. **(B)** Box and whisker plot showing that loci with significant gain or reduction of Ctcf binding in *Eed*^-/-^ cells also have corresponding significant change in Cdx2 binding. **(C)** Gene ontology analysis for genes within 50 kb of the loci with alterations in both Ctcf and Cdx2 binding as compared to genes similarly linked to loci with change in only Ctcf binding. **(D)** Representative genomic tracks showing relatively stable histone modifications at locus with Cdx2 gain in adult *Eed*^-/-^ cells; PRC2 mediated H3K27 methylations are lost upon induced deletion of *Eed*.

